# scBubbletree: Quantitative visualization of single cell RNA-seq data

**DOI:** 10.1101/2023.03.09.531263

**Authors:** Simo Kitanovski, Yingying Cao, Dimitris Ttoouli, Farnoush Farahpour, Jun Wang, Daniel Hoffmann

## Abstract

**Motivation:** Visualization approaches transform high-dimensional data from single cell RNA sequencing (scRNA-seq) experiments into two-dimensional plots that are used for analysis of cell relationships, and as a means of reporting biological insights. Yet, many standard approaches generate visuals that suffer from overplotting, lack of quantitative information, and distort global and local properties of biological patterns relative to the original high-dimensional space.

**Results:** We present scBubbletree, a new, scalable method for visualization of scRNA-seq data. The method identifies clusters of cells of similar transcriptomes and visualizes such clusters as “bubbles” at the tips of dendrograms (bubble trees), corresponding to quantitative summaries of cluster properties and relationships. scBubbletree stacks bubble trees with further cluster-associated information in a visually easily accessible way, thus facilitating quantitative assessment and biological interpretation of scRNA-seq data.

**Availability and Implementation:** the R package scBubbletree is freely available at: https://bioconductor.org/packages/scBubbletree/

## 1 Introduction

Single cell RNA sequencing (scRNA-seq) data can convey an unprecedented richness of biological information, which has led to an explosion of scRNA-seq experiments [Svensson et al., 2018]. However, the complexity of scRNA-seq data makes analysis also notoriously difficult: the transcriptome of each cell is characterized by a high-dimensional vector of gene expressions, and we have many cells and hence many vectors. How can we visually present such complex data so that essential biological information becomes easily accessible? Standard workflows perform reduction of the high-dimensional transcription data into printable two dimensional scatterplots [McInnes et al., 2018, Van der Maaten and Hinton, 2008] with each cell corresponding to one dot and the geometric arrangement of the dots reflecting aspects of transcriptional similarity of cells. While this approach works reasonably well for small samples, visualization of large and complex samples runs into the well-known problem of overplotting [Carr et al., 1987], i.e. massive piling up of dots on top of each other severely hampers user’s ability to identify patterns in the plotted data. This issue is compounded by distortions in the sample’s distance structure [Huang et al., 2022] that arise as we squeeze high-dimensional transcription data into two dimensions. Thus, visuals produced with the standard workflow are in general inept for presenting quantitative information, including also transcriptional similarity.

Here we present scBubbletree, a method for quantitative visual exploration of scRNA-seq data. When designing scBubbletree our guiding question was: Which properties of scRNA-seq data should be visualized? We identified three classes of properties: i) local and global transcriptional structure, ii) cell density distribution and iii) cell attributes (marker gene expressions, biological condition, cell type labels, etc.). Furthermore, scalability was an important factor in the design of scBubbletree, i.e. our method was developed to avoid overplotting. These goals were achieved by integrating in scBubbletree state-of-the-art methods for clustering of scRNA-seq data and for visualization. scBubbletree is available as an R-package that is open-source, easy-to-use and simple to integrate with popular approaches for scRNA-seq data analysis.

We demonstrate the added value of visual exploration with scBubbletree compared to popular methods, such as UMAP and t-SNE, by analyzing two public datasets.

## 2 Materials and methods

scBubbletree is implemented as an R package that provides an easy-to-use workflow for visual exploration of single cell RNA-seq data. A typical application, as presented here with two scRNA-seq samples (studies A and B), runs in less than one hour on a standard personal computer. Furthermore, scBubbletree can be integrated seamlessly with the R package Seurat [Hao et al., 2021], a popular tool for scRNA-seq analysis. In the next we describe the implementation of scBubbletree in detail.

### 2.1 Input

The pipeline takes as first input a matrix *A*^*n×f*^ that represents a low-dimensional projection of the high-dimensional scRNA-seq data, with *n* rows as cells and *f* columns as low-dimensional features. scBubbletree works directly with *A*^*n×f*^ and is agnostic about the initial data processing protocol. For the generation of the low-dimensional projection, we recommend the use of linear techniques that approximately conserve pairwise distances between cellular transcriptome vectors. Example techniques are principal component analysis (PCA) [Hotelling, 1933], non-negative matrix factorization [Paatero and Tapper, 1994], or multi-dimensional scaling [Kruskal, 1964]. *Non-linear* dimensionality reduction techniques, such as t-SNE [Van der Maaten and Hinton, 2008] and UMAP [McInnes et al., 2018], should be avoided as these approaches tend to distort long-range distances [Becht et al., 2019]. In the present work we used PCA for the low-dimensional projection.

### 2.2 Algorithm

In short, the algorithm of scBubbletree identifies clusters of transcriptionally similar cells, and then visualizes these clusters as leaf-nodes (bubbles) of a hierarchical dendrogram (bubbletree). The workflow comprises four steps (Fig. 1A): 1. determining the clustering resolution, 2. clustering, 3. hierarchical cluster grouping, and 4. visualization. We explain each step in the following.

**Figure 1:**
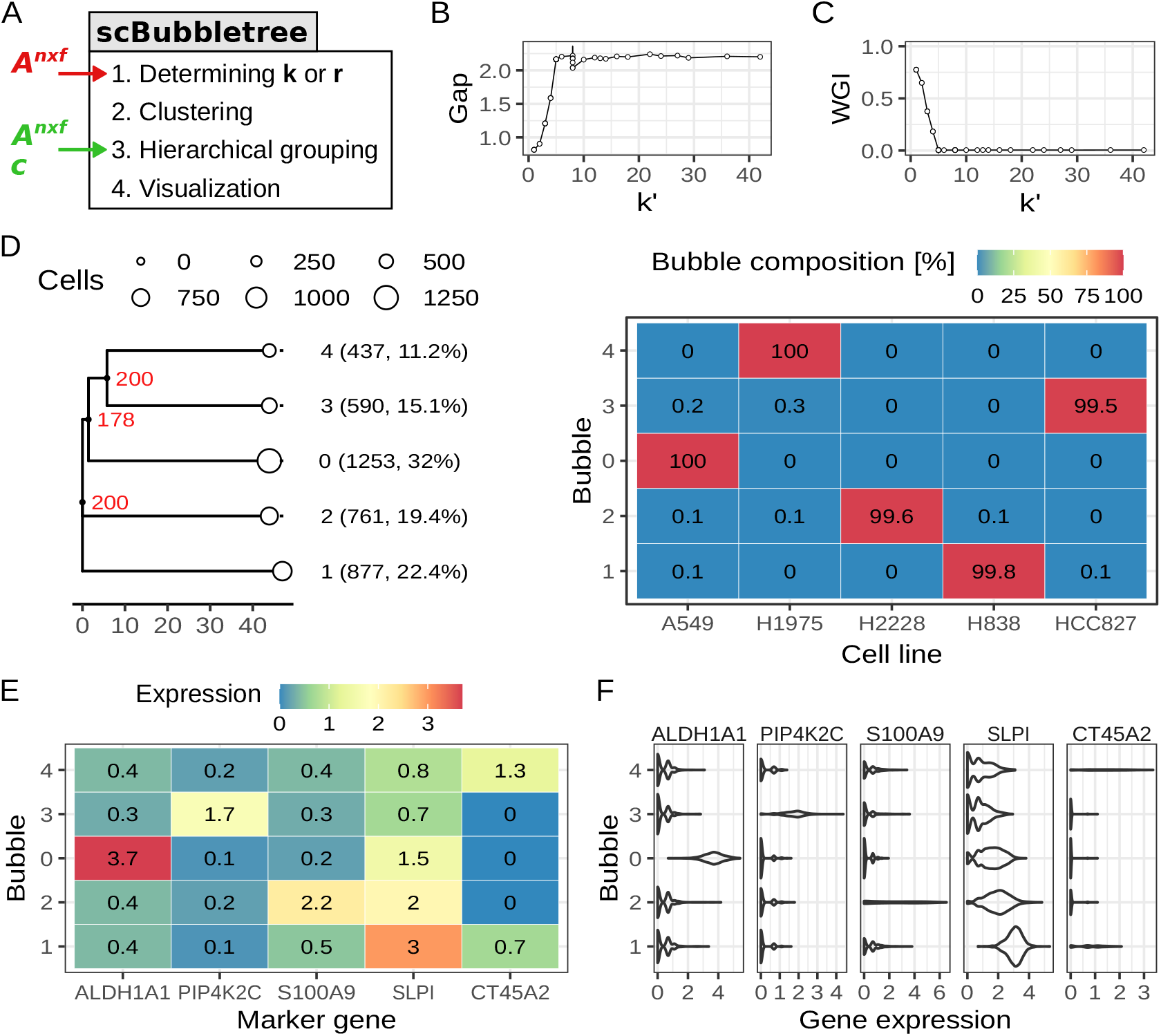
scBubbletree analysis of dataset A. (A) scBubbletree workflow with main and alternative inputs shown with red and green labels, respectively. (B) Gap statistic for clustering solutions generated by the Louvain method with varying clustering resolutions. Vertical error bars are 95% confidence intervals of the gap statistic. (C) WGI for clustering solutions in panel B and the predicted cell lines of dataset A. (D) bubbletree (tree structure, left) annotated with a heatmap (right) displaying as rows the within-bubble relative frequencies of different cell lines. The bubbletree has five bubbles (white points) shown as leaves. Bubble radii scale linearly with the number of cells in the bubbles. Bubble identities, absolute and relative cell frequencies are shown as labels between bubbletree and heatmap. Red labels are branch bootstrap support values. Rows in the heatmap integrate to 100%. (E) Mean normalized expression of five marker genes (x-axis) in each bubble (y-axis). (F) Density distribution of five marker gene (column panels) with normalized expressions of the cells in each bubble shown as violins. Numbers in heatmap tiles are rounded to the nearest tenth.

#### 2.2.1 Determining the clustering resolution with gap statistic

scBubbletree can cluster scRNA-seq data in two ways, namely by graph-based community detection (GCD) algorithms such as Louvain [Blondel et al., 2008] or Leiden [Traag et al., 2019], and by k-means [Macueen et al., 1967]. Benchmarking studies have demonstrated that approaches for GCD, such as Louvain, perform well with scRNA-seq data and have relatively short runtimes [Duò et al., 2018]. These findings were independently verified by our benchmarking study of the impact of clustering algorithm on biological cluster homogeneity (Supplementary Section 1). We recommend using the original Louvain algorithm for clustering of large scRNA-seq datasets. For smaller and simpler scRNA-seq datasets k-means generates comparable results.

Each clustering approach requires a resolution as input. In k-means, the clustering resolution is specified by the parameter *k* (*k* ∈ ℕ) that determines the number of clusters in the data, whereas Louvain (or Leiden) uses a resolution parameter *r* (*r* ∈ ℝ_>0_), where higher *r* lead to more clusters and lower *r* lead to fewer clusters. Hence, *r* can be mapped onto a number *k′* of communities, with a range of *r* values of mapping onto the same value of *k′*. The choice of *k* and *r* should be guided primarily by the sample heterogeneity: the higher the cellular diversity in the sample, the higher the clustering resolution needed to resolve that diversity. The choice of *k* and *r* is also impacted by the research objective, for instance, high clustering resolution might be necessary to identify e.g. rare T cell subsets in a large sample of immune cells, whereas lower clustering resolution might be sufficient to characterize canonical immune subsets of that sample.

To determine *k* and *r*, scBubbletree relies on the *gap statistic* method [Tibshirani et al., 2001] as implemented in the R-package cluster (function *clusGap*, version 2.1.2). This functionality is implemented in scBubbletree’s functions *get_k* and *get_r*. The gap statistic compares the within cluster sum of squares (WCSS) to its expectation under an appropriate null reference distribution for different values of *k* and *r*. To estimate *k* and *r* we need to examine the gap curve for a) clear maximum or b) an ‘elbow’ (a bend in the curve from high to low slope). The functions *get_k* and *get_r* report the average gap and its 95% confidence interval for each clustering resolution computed based on *B*_gap_ simulations.

Many complementary approaches for determining the clustering resolution are available in the literature [Yu et al., 2022], and these can be used alongside the gap statistic. Furthermore, we recommend that the estimated *k′* (associated with *r*) and *k* be compared against prior biological knowledge about the cellular composition based on data from e.g. the human protein atlas (https://www.proteinatlas.org) or from relevant publications.

### 2.2.2 Determining the clustering resolution with Gini impurity

For some scRNA-seq datasets it is possible to assess the subtypes of individual cells by mapping their transcriptional profiles onto reference atlases [Aran et al., 2019]. scBubbletree quantifies the homogeneity (or ‘purity’) of individual clusters *i* of *n*_*i*_ cells in terms of the their composition of subtype labels with the Gini impurity (GI) index:

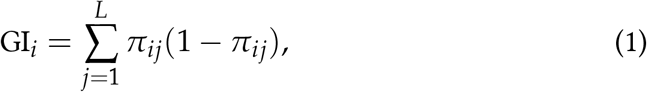

where *L* is the total number of subtype labels, and *π*_*ij*_ the relative frequency of label *j* in cluster *i*. In homogeneous clusters, GI takes on a small value close to zero, and in mixed clusters GI can approach 1 − (1/*L*). Given a clustering result with *k* clusters, *get_gini* also computes the weighted Gini impurity (WGI) index by calculating the weighted average of the GIs:

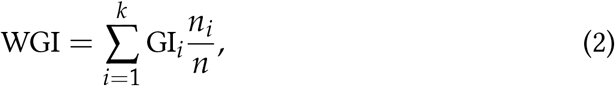

with *n* = ∑_*i*_ *n*_*i*_. We expect low WGIs for clustering results at optimal clustering resolutions, and higher WGIs for clustering results at insufficient resolutions. Hence, the WGI can be used like the gap statistic to identify appropriate resolutions.

#### 2.2.3 Clustering

For clustering, scBubbletree uses as main inputs matrix *A*^*n×f*^, a clustering algorithm, and the clustering resolution (*k* or *r*) estimated from the previous step of this workflow. Four algorithms for GCD are available via the function *get_bubbletree_graph*, including, the original Louvain algorithm [Blondel et al., 2008], the Louvain algorithm with multilevel refinement (LMR) [Rotta and Noack, 2011], the smart local moving (SLM) algorithm [Waltman and Van Eck, 2013], and the Leiden algorithm [Traag et al., 2019]. In addition to this, k-means [Macueen et al., 1967] clustering can be performed with the function *get_bubbletree_kmeans*. Auxiliary input parameters can be used to employ a specific variant of the clustering algorithms and to fine-tune their performance. To perform GCD, scBubbletree first constructs a shared nearest neighbor (SNN) graph using the function *FindNeighbors* (R-package Seurat, version 4.1.0). GCD is applied with Seurat’s function *FindClusters* and k-means clustering by the function *kmeans* (R-package stats, version 4.2).

The output of the clustering step is a vector *c* = (*c*_1_, *c*_2_, …, *c*_*n*_) of cluster assignments for the *n* cells, with *c*_*i*_ ∈ *{*1, 2,…, *k}*. The clustering result allows us to partition *A*^*n×f*^ into *k* data subsets 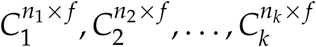, with *n*_*i*_ as the number of cells in cluster *i* and *n* = ∑_*i*_ *n*_*i*_. These matrices are used to compute inter-cluster distances in the next step of this workflow.

To facilitate the use of novel clustering approaches [Kiselev et al., 2019], scBubbletree provides the function *get_bubbletree_dummy*, which allows the user to provide as input matrix *A*^*n×f*^ and vector *c* of cluster assignments (Fig. 1A, green input) and proceed with hierarchical grouping of the clusters. This enables seamless integration of scBubbletree with computational pipelines that employ other clustering approaches, such as PhenoGraph [Levine et al., 2015] or TooManyCells [Schwartz et al., 2020].

#### 2.2.4 Hierarchical grouping of clusters

Clusters of cells are arranged in a dendrogram by performing agglomerative hierarchical clustering with average linkage [Hastie et al., 2009] (function *hclust*, R-package stats, version 4.2). Starting at the lowest level with *k* clusters (or communities) of cells, the clustering procedure selects pairs of clusters with the smallest inter-cluster Euclidean distance and groups them at the next higher level of the hierarchy. This is repeated until a complete dendrogram is built with only one cluster at the highest level that contains the full data. Other commonly used linkage functions and the Manhattan distance metric are also implemented in scBubbletree.

The distance between clusters *i* and *j* can be computed by estimating the average Euclidean distance between all pairs of cells in 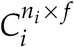 and 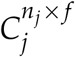. Hence, the complete hierarchical clustering procedure has time complexity *O*(*n*^2^) and requires approximately Ω(*n*^2^) memory to store the resulting distance matrix. Assuming that each matrix cell takes up 8 bytes of memory, for a dataset of 10^6^ cells this operation requires 7,450 gigabytes of memory, and considerable computational cost.

To avoid such high memory demands scBubbletree employs a bootstrapping approach with *B* iterations (Supplementary Algorithm 1). In iteration *b*, the algorithm draws with replacement a random subset with 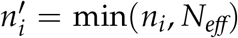 and 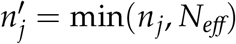 number of cells from cluster *i* and *j*, respectively, where *N*_*eff*_ is a user-defined parameter that controls the maximum number of cells to be drawn from each cluster. The algorithm computes the average distance between cluster *i* and *j* based on 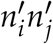 inter-cell Euclidean distances, and stores the result in distance matrix 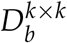:

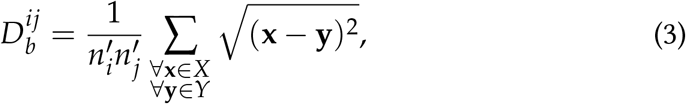

where 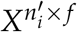 and 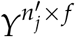 are subsets drawn from 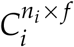 and 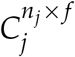, respectively, and **x** and **y** are cell transcriptome features (row vectors) from each matrix. At the end of bootstrap iteration *b*, the distance matrix 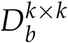 is provided as input for hierarchical clustering with average linkage to generate the dendrogram *H*_*b*_.

From the collection of *B* distance matrices we compute an average distance matrix 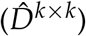, and use 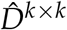 to generate a consensus hierarchical dendrogram (bubbletree; *Ĥ*) by hierarchical clustering with average linkage. The collection of bootstrap dendrograms are used to assess the robustness of the bubbletree topology by quantifying the number of times each branch in *Ĥ* was found among the topologies of the bootstrap dendrograms (function *prop*.*clades*, R-package ape, version 5.6.2). Branches can have variable degrees of support ranging between values close to 0 (no support) and *B* (complete support). Distances between individual bubbles are visualized quantitatively as sums of branch lengths in root-to-leaf direction between these bubbles.

#### 2.2.5 Visualization of the bubbletree

The main visual output of scBubbletree is a bubbletree (Fig. 1D and Fig. 2C). The bubbles represent clusters of cells of similar transcriptomes, and the tree describes hierarchical relationships and distances between the bubbles. Bubbletrees are visualized with ggtree (R-package, version 3.2.1), which offers a rich syntax for visualization of dendrograms [Yu, 2020]. ggtree extends the flexible ggplot2 (R-package, version 3.3.5) visualization framework [Wickham, 2016], and thereby allows multiple layers of annotations to be attached to the bubbletree.

**Figure 2:**
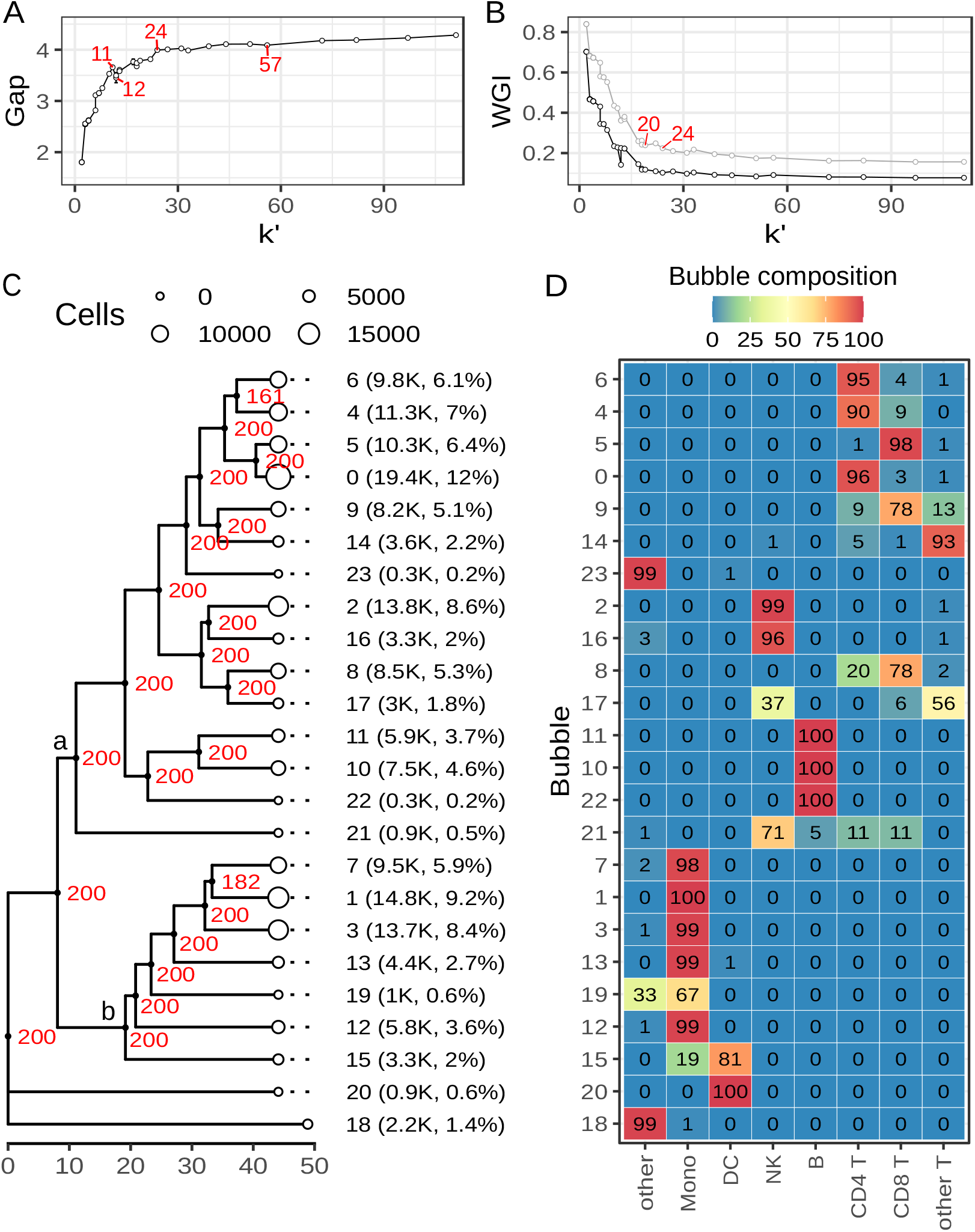
scBubbletree analysis of dataset B. (A) Gap statistic for clustering solutions generated by Louvain clustering algorithm with varying clustering resolutions. Vertical error bars are 95% confidence intervals of the gap statistic. (B) WGI index for clustering solutions by Louvain algorithm and the cell type annotation sets *l1* (gray points and lines) and *l2* (black points and lines). (C) bubbletree with 24 bubbles (white points) shown as leaves. Bubble radii scale linearly with the number of cells in the bubbles. Bubble identities, absolute (in thousands) and relative cell frequencies are shown as labels. Red branch labels are branch bootstrap support values. Clades of lymphocytes and monocytes are labeled with symbols ‘a’ and ‘b’. (D) Within-bubble relative frequencies (%) of cell types (*l1*; x-axis) rounded to the nearest integer. Rows integrate to 100% (up to rounding error).

scBubbletree scales the bubble radii linearly as the number of cells in the corresponding bubbles. As linear increase in bubble radii leads to quadratic increase in bubble areas, this scaling visually emphasizes bubbles with higher numbers of cells. Each bubble is labelled with an identifier and its absolute and relative numbers of cells.

In summary, a bubbletree gives a quantitative visual summary of scRNA-seq data in terms of transcriptome cluster sizes and relationships.

#### 2.2.6 Visualization of cell attributes

To make bubbles biologically interpretable, bubbletrees facilitate addition of further information, including *numeric* cell attributes, such as expression levels of marker genes, number of unique molecular identifiers (UMIs), or prevalence of UMIs from mitochondrial genes; or *categorical* cell attributes, such as predicted subtype labels, treatment groups (cancer vs. control cell), sample name, or cell cycle phase. For visualization of numeric and categorical cell attributes scBubbletree relies on the R-package ggplot2. Assembly of plots from multiple cell attributes is done with the R-package patchwork (version 1.1.1), which enables attaching of user-generated ggplot2 figures to the bubbletree.

scBubbletree provides two functions for visualization of numeric cell attributes: the function *get_num_tiles* computes statistical summaries (mean, median, sum, number of zero/nonzero values, etc.) of numeric cell attributes in each bubble and visualizes them with tile plots; the function *get_num_violins* visualizes the distributions of numeric cell attributes in each bubble with violin plots.

Categorical cell attributes are visualized using a matrix of tiles in which columns represent specific attribute categories. Tile colors and annotations can be configured for two types of interpretation using the logical parameter *integrate_vertical*. For *integrate_vertical=T* the tiles will show the relative frequencies of each attribute label across the different bubbles, which answers questions such as: are cells carrying a specific label (e.g. ‘T cell’ or sample label ‘S1’) enriched in a particular set of bubbles? Alternatively, for *integrate_vertical=F* the tiles will show the within-bubble relative frequencies of different labels. With this information we can answer questions such as: what is the label composition in a particular bubble?

### 2.3 Evaluation of scBubbletree using publicly available data

scBubbletree was evaluated with two publicly available scRNA-seq datasets, which we call A and B; for detailed descriptions of datasets and workflow see Supplementary Section 2.

Dataset A contains 3,918 cell transcriptomes from five human lung adenocarcinoma cell lines (HCC827, H1975, A549, H838 and H2228) [Tian et al., 2019]. The predicted cell line labels for each cell are available as part of the meta data. Dataset B contains transcriptomes of 161,764 peripheral blood mononuclear cells (PBMCs) from eight healthy volunteers enrolled in an HIV vaccine trial [Hao et al., 2021]. PBMC subtype predictions are available for each cell at two levels of resolution: annotation set *l1* and *l2* with 8 and 31 subtype categories, respectively.

In both datasets we saw that the first 15 principal components capture most of the variance in the data, and the proportion of variance explained by each subsequent principal component was negligible (Supplementary Fig. S1A-B). Thus, we used the single cell projections in 15-dimensional feature space, *A*^3,918*×*15^ and *A*^161,764*×*15^, as input of scBubbletree.

## 3 Results

### 3.1 Study A: Exploring sample of five cancer cell lines

#### 3.1.1 Generating the bubbletree

We know that dataset A contains a mixture of five cancer cell lines. Hence, we expect *k* = 5 cell clusters or communities. To verify this we computed the gap statistic for the Louvain method with resolution parameter *r* = 10^*R*^, where *R* was initialized using a sequence of values from -4 to 1 in increments of 0.1. At each resolution *r* we recorded the number *k′* of identified communities (Supplementary Fig. S2). WGIs were computed based on the cluster assignments and the vector of predicted cell line labels.

The gap curve had a distinct ‘elbow’ at *k′* = 5 (*r* ∈ [0.0025, 0.12]) (Fig. 1B). In line with these observations, we saw a steep drop in the WGIs to a value close to 0 at *k′* = 5 (*r* = 0.0025) (Fig. 1C) These results suggested that there is little benefit in using clustering resolutions that yield more than five communities.

In the next step of the scBubbletree workflow we applied clustering with the original version of Louvain [Blondel et al., 2008] (function *get_bubbletree_graph*) and resolution parameter *r* = 0.0025 (*k′* = 5). For input, we constructed a SNN graph based on *A*^3,918*×*15^, where each vertex (cell) was connected to its 50 nearest neighbors. Clustering was performed with 20 random starts and up to 100 iterations in each run. The clusters were organized in a bubbletree by hierarchical clustering with average linkage (tree in Fig. 1D). We ran *B* = 200 bootstrap iterations and drew samples with up to *N*_*eff*_ = 200 cells from each cluster to estimate inter-cluster distances and their robustness.

#### 3.1.2 Examining the bubbles

Bubble sizes range from the smallest (bubble 4, 437 cells, 11.2% of the sample cells) to the largest (bubble 0, 1,253 cells, 32% of the sample cells). By visualizing the cell lines as categorical attributes, we saw clear mapping between different cell lines and bubbles (heatmap in right part of Fig. 1D), i.e. cells from a specific cell lines were enriched in specific bubbles. For instance, 100% of the cells in bubble 0 and 4 belonged to cell line A549 and H1975, respectively, and between 99.5% and 99.8% of the cells in bubbles 1, 2 and 3 belong to cell lines H838, H2228 and HC827, respectively.

By visualizing in each bubble the mean (normalized) expression (Fig. 1E) and the expression distributions (Fig. 1F) of five marker genes (one per cell line), we were able to confirm that bubble 0 was not only enriched with cells from cell line A549, but also had high ALDH1A1 expression which is a known A549 marker [Park et al., 2017]. Similarly, bubble 4 had high expression of CT45A2 which is a marker of H1975 [Yang et al., 2019]. Bubbles 1, 2, and 3 were associated with high expression of SLPI, S100A9, and PIP4K2C, respectively, and based on data from the Cancer Dependency Map Portal (https://depmap.org; accessed 23.06.2022) we were able to confirm that these genes are typically over-expressed in cell lines H838, H2228, and HCC827, respectively.

#### 3.1.3 Examining the tree

The distance between bubble 3 and 4 was smaller than that of any other pair of bubbles in the bubbletree (Fig. 1D). This hinted at relatively higher transcriptional similarity between the corresponding cell lines HCC827 and H1975. Furthermore, bubble 3 and 4 were in a robust sub-tree that was found in all 200 bootstrap dendrograms. The sub-tree of bubbles 0, 3, and 4 was less robust (178 of 200 bootstrap dendrograms).

The robust sub-tree of bubbles 3 (cell line HCC827) and 4 (cell line H1975) with the relatively short distance between these bubbles indicates transcriptional similarity of these cell lines. We checked this aspect of the tree by comparison with a set of independently measured transcriptional profiles of 69 adenocarcinoma cell lines from the Cancer Cell Line Encyclopedia [Barretina et al., 2012], including HCC827 and H1975 (Supplementary Section 3). Euclidean distances were computed between the vectors of normalized gene expressions of all pairs of cell lines. The distance between HCC827 and H1975 was the 6th lowest among 2,346 pairwise comparisons (Supplementary Fig. S3), whereas the distances between the remaining cell lines of dataset A fell well within the general distribution of distances between the different adenocarcinoma cell lines. This result is consistent with the structure of the bubbletree, especially the robust closeness of bubbles 3 and 4.

Dataset A had been measured with an artificially mixture of five distinct cell line populations, which explains the simple structure of the resulting bubbletree. The recovery of the ground truth of five cell lines (Fig. 1), was a sanity check for scBubbletree. In this simple case, even UMAP and t-SNE were able recover aspects of the overall structure (Supplementary Fig. S6A,B). The advantage of bubbletrees over these standard methods was the quantitative visualization of tree structure and cluster sizes. However, scBubbletree has been developed for the analysis of more complex scRNA-seq data that are typically observed with real tissue samples, as in the following study B.

### 3.2 Study B: Exploring a sample of human PBMCs

#### 3.2.1 Clustering resolution

To determine the clustering resolution of dataset B we computed the gap statistic for the Louvain method with resolution parameter *r* = 10^*R*^, where *R* was initialized using a sequence of values from -4 to 1 in steps of 0.1 (Fig. 2A). At each resolution we recorded the number *k′* of identified communities (Supplementary Fig. S4).

The gap increased rapidly between *k′* = 1 and *k′* = 11, followed by a dip at *k′* = 12 and a further ascend to about 4 at *k′* = 24 (*r* ≈ 0.79) (Fig. 2A), and a much slower, approximately linear increase at higher *k′*. The increase in the gap between *k′* = 22 and *k′* = 24 was about two-fold larger than that in the interval between *k′* = 24 and *k′* = 57, even though only two communities were added in the first interval and 23 communities (about 10-fold more) were added in the second interval. This suggested that there is little benefit in using values of *r* larger than 0.79 (*k′* = 24).

The value of *k′* = 24 is close to the number of 18 canonical cell populations identified in PBMCs in the Human Protein Atlas (HPA) database (https://www.proteinatlas.org/humanproteome/immune+cell).

#### 3.2.2 Clustering resolution analysis based on WGI

To compute WGI curves we used as inputs the PBMC subtype labels from annotation set *l1* and *l2* and the Louvain clustering assignments generated for the gap analysis. We saw that annotation set *l1* (Fig. 2B) was associated with lower WGIs compared to *l2*. The relatively lower WGIs of *l1* are likely due to the fact that *l1* contains labels of 8 major PBMC subtypes that have distinct transcriptional profiles. Hence, in each bubble of dataset B we expect a low degree of mixing in terms of different *l1* labels (Fig. 2D). In contrast to this, annotation set *l2* contains labels of 31 PBMC subtypes including, for instance, 13 T cell subtypes with comparable transcriptional profiles. Hence, we expect relatively higher degree of mixing of different *l2* labels in common clusters (heatmap in Supplementary Fig. S5A) in comparison to the *l1* labels.

The WGI curve converged to low WGIs at *k′* ≈ 20 for both annotation sets. This is similar to the clustering resolution of *k′* = 24 identified with the gap curve.

#### 3.2.3 Clustering and generating the bubbletree

For clustering of dataset B we chose the original Louvain method and a resolution parameter *r* = 0.79 (*k′* = 24). First, we constructed a SNN graph based on *A*^161,764*×*15^, where each vertex (cell) in the SNN graph was connected to its 50 nearest neighbors. Clustering was performed with 20 random starts and up to 100 iterations in each run. The clusters were organized in a bubbletree using hierarchical clustering with average linkage (Fig. 2C). For this we used *B* = 200 bootstrap iterations and drew samples with up to *N*_*eff*_ = 200 cells from each cluster to compute inter-cluster distances.

The resulting bubbletree (Fig. 2C) had 24 bubbles with sizes ranging from 19,400 cells (12%, cluster 0) down to 300 cells (0.3%, cluster 23). The bubbletree had two major clades and two small outgroup bubbles (Fig. 2C). Clade ‘a’ contained 15 bubbles that accounted for about 65% of the cells in the sample, whereas clade ‘b’ contained seven bubbles that accounted for about 32% of the cells in the sample. Nearly all branches in the bubbletree were completely robust. The branches between bubble 4 and 6 and between bubble 1 and 7 had lower branch support as they were found in only 161 (80.5%) and 182 (91%) out of 200 bootstrap dendrograms, respectively. A biological interpretation of the different clades and their within-clade branching patterns are provided in the following.

#### 3.2.4 Bubbletree evaluation with annotation set *l1*

To determine which PBMC subtypes are enriched in each bubble we visualized the within-bubble relative frequencies of 8 major PBMC subtypes from annotation set *l1* (heatmap in Fig. 2D). Note that Figs. 2C and 2D demonstrate that bubbletrees (Fig. 2C) can be easily and coherently stacked with additional quantitative information (Fig. 2D), thus facilitating biological interpretation (see Supplementary Fig. S5 for more extensive stacking).

Fig. 2D shows that most bubbles are enriched with cells from either one PBMC subtype or a combination of related subtypes (e.g. CD4^+^ and CD8^+^ T cells). Furthermore, bubbles enriched with cells from related PBMC subtypes are close to each other in the bubbletree; for instance, bubbles 10, 11, and 22 are enriched with B cells and constitute a robust subclade of the bubbletree. Similarly, bubbles 7, 1, 3, 13, 19, 12 and 15 form clade ‘b’ and are enriched with monocytes and dendritic cells (DCs).

The bubbles of clade ‘a’ are mostly enriched with lymphocytes, and robust subclades within clade ‘a’ are formed by sets of bubbles that are enriched with specific lymphocyte subtypes, including T cells, B cells, and NK cells. The small outgroup bubble 21 contains a mixture of different lymphocytes. Clade ‘b’ has seven bubbles that are enriched in monocytes and DCs. The branches in that clade form a ladder-like topology in contrast to the more complex nested topology of clade ‘a’. The simpler topology in ‘a’ may be due to the lower heterogeneity of circulating monocytes compared to lymphocytes, with classical monocytes constituting about 85% of all circulating monocyte pool in humans [Patel et al., 2017].

Clearly separated from clades ‘a’ an ‘b’ is the small outgroup of bubbles 20 of DCs and 18 consisting of a mixture of cells (‘other’).

Supplementary Section 4 contains a bubbletree analysis based on the more detailed annotation set *l2*, revealing enrichment of specific PMBC subtypes in the different bubbles.

### 3.3 Comparison with known approaches

We compared scBubbletree with UMAP, t-SNE and other popular methods for scRNA-seq data visualization with respect to visualization clarity, conservation of local and global data structure.

#### 3.3.1 Conservation of local structure: qualitative analysis

In the following we compare the ability of the Louvain method to preserve local structure with those of UMAP and t-SNE. For this analysis we generated conventional UMAP and t-SNE figures for datasets A and B (Supplementary Fig. S6A-D). In each figure, the different bubbles in dataset A and B were represented by heavily overplotted groups of dots in 2D UMAP and t-SNE space so that group size cannot be discerned from the figure. To our surprise, some cell lines from dataset A were split by UMAP and t-SNE into multiple subclusters; for instance, the cell line H1975 (bubble 4) had three distinct subclusters. The apparent distances between these subclusters in the drawing plane was more pronounced in t-SNE space (Supplementary Fig. S6B). Similarly, cell lines A549 (bubble 0) and H838 (bubble 1) were split into at least two subclusters of cells (Supplementary Fig. S6A-B). We realized that the cells from the smaller subclusters of A549 and H838 had systematically lower numbers of detected RNA molecules compared to the cells from the larger subclusters (Supplementary Fig. S7C-F). In contrast to this, the three subclusters in H1975 had comparable numbers of detected RNAs (Supplementary Fig. 7A-B), i.e. the subcluster structure of cell line H1975 could not be explained by differences in the number of detected RNA molecules. By using higher clustering resolution as input of Louvain we recovered these subclusters (data not shown). However, this was associated with negligible improvement in the gap and WGI (Fig. 1B-C).

Several conclusions can be drawn from this qualitative analysis. First, visualization of the local structure of scRNA-seq data with UMAP and t-SNE can be challenging even for relatively small samples. Second, UMAP and t-SNE are prone to generating clearly separated visual clusters despite few underlying biological differences [Huang et al., 2022]. Furthermore, UMAP and t-SNE use a number of hyperparameters, which can have severe impact on the 2D projection [Huang et al., 2022, McInnes et al., 2018] (Supplementary Fig. S8). Taken together, it is difficult to ascertain whether the subclusters in dataset A represent biologically distinct subpopulations of the main cell lines or technical artifacts resulting from biological/technical noise.

#### 3.3.2 Preservation of distance structure

Users of conventional methods such as UMAP or t-SNE are tempted to interpret distances between dots or apparent clusters as indicators of transcriptional similarity of the corresponding cells. Yet, several studies [Kobak and Linderman, 2021, McInnes et al., 2018] have warned that while both, UMAP and t-SNE, consistently preserve distances on a small-scale, accurate preservation of large-scale distances is not always guaranteed.

To highlight the challenge in preserving large-scale distances with conventional methods, we analyzed UMAP and t-SNE plots of a dataset with over 1.3 million mouse brain cells (Supplementary Fig. S9A-B). The UMAP and t-SNE plots suffer from excessive overplotting, which hinders visual evaluation of the distances between cells and clusters. In addition to this, we see large spread (indicated by the wide and overlapping error bars in Supplementary Fig. S9C-D) in the distances between cells in 20D PCA space over the entire range of distances in 2D UMAP and t-SNE space, which indicates ambiguous mapping of distances in 2D UMAP and t-SNE space relative to distances in 20D PCA space. We saw similar patterns in the 2D UMAP and t-SNE plots of dataset B (Supplementary Fig. S6G-H), even though dataset B is about one order of magnitude smaller than the dataset of mouse brain cells. As currently available workflows for scRNA-seq are able to generate data at similar scale and complexity as in these examples, we conclude that inference of transcriptional relationships between cells and clusters from UMAP and t-SNE will in general not be reliable.

Bubbletrees by construction should better reflect distances in PCA space in the dendrogram distances. In fact, Supplementary Figs. S6I-J and 10B demonstrate that bubbletree distances are consistent with the average distances in PCA space between cells from the corresponding bubbles. This was true for distances on all scales except for intra-bubble distances, which correspond to bubbletree distance = 0 (distances between bubbles to themselves). For bubbletree distances equal to 0 we saw consistently small average distances between cells in PCA space. It is noteworthy that the spread of distances between dots in PCA space was not uniform over all pairs of bubbles, presumably because the underlying clusters deviate more or less from homogeneous spherical shapes in PCA space, leading to the dispersion of inter-bubble cell distances.

#### 3.3.3 Visualization clarity

We assessed the visualization clarity of scBubbletree with respect to two aspects: 1) *scalability*: the ability to handle large RNA-seq datasets; 2) *flexibility*: the ease of incorporating diverse cell attributes as part of the main visuals.

##### Scalability

We investigated scalability in the context of the overplotting problem. A high degree of overplotting was observed in the 2D UMAP and t-SNE maps generated based on dataset A and B (Supplementary Fig. S6A-D). The overplotting was particularly excessive in the 2D UMAP and t-SNE maps of dataset B, where individual dots (cells) were barely visible and clusters of cells appeared as colored blobs. There are several consequences of such massive overplotting: 1. failure to grasp data’s local and global structure; 2. inadequate description of cluster homogeneity in terms of different cell attributes; 3. loss of information about the density of cells across the different clusters; and 4. visual biases, i.e. users can deliberately or inadvertently emphasize or de-emphasize specific cell attributes by assigning the corresponding cells to the top of the cell piles [Freytag and Lister, 2020].

Various techniques have been proposed to mitigate the overplotting problem, such as, reducing the size of the dots, adding an alpha channel for point transparency, data subsetting [Hao et al., 2022], hexagonal cell binning [Freytag and Lister, 2020], applying density-preserving dimensionality reduction [Narayan et al., 2021] or exploring the data with interactive graphical tools [Hillje et al., 2020]. However, as the scale of scRNA-seq datasets continues to increase these countermeasures are of limited help, and most of them even have a negative impact on the clarity of the resulting figures.

Instead of visualizing individual cells, scBubbletree uses a high-level abstraction (bubbletree) to summarize scRNA-seq datasets and thereby avoids the overplotting problem. To demonstrate the high degree of scalability of scBubbletree in comparison to UMAP and t-SNE, we visualized with each approach a dataset of over 1.3 million mouse brain cells (Supplementary Fig. S9A-B and S10A). While both visuals are complex, the bubbletree approach at least offers an opportunity to understand the dataset, whereas the UMAP and t-SNE visuals lump all cell clusters into densely packed areas, making it impossible to understand the structure of the data. scBubbletree is not the only approach that employs trees to visualize scRNA-seq data, i.e. approaches such as TooManyCells [Schwartz et al., 2020] and clustree [Zappia and Oshlack, 2018] use similar strategies to mitigate the overplotting problem.

##### Flexibility

The flexibility of scBubbletree was compared with that of UMAP, t-SNE, TooManyCells and clustree. Specifically, we assessed the ease with which each tool is able to integrate and combine multiple cell attributes.

Cell attributes are visualized in UMAP and t-SNE plots by color-coding each point according the value of a particular attribute (e.g. expression level of a marker gene). For simultaneous visualization of two or more cell attributes, it is current practice to generate multiple and often small copies of the UMAP or t-SNE plots in which the individual attributes are visualized [Stuart et al., 2019]. Although this method of integrating and presenting scRNA-seq data is widely used, it suffers from the same and even more shortcomings than individual UMAP and t-SNE plots.

TooManyCells is more flexible as it uses multiple facets to visualize cell attributes alongside its dendrogram e.g. by adjusting the color-code or the widths of its branches to for visual summaries (e.g. averages) of continuous cell attributes. TooManyCells can also attach pie charts to each leaf-node to visualize the within-leaf composition of a categorical attribute. Similarly, clustree visualizes cell attributes as part of its main dendrogram by adjusting the different visual elements such as colors, sizes, shapes and labels of the leaf-nodes. Plots that encode information in such a diverse set of visual elements make it difficult for the viewer to decode and discover relationships. With scBubbletree we have therefore limited the number of types of visual elements (dendrogram, bubbles, and annotations) but made it easy to attach to the tree simple, matrix-like plots (heatmaps, violins) with one matrix row for each bubble (e.g. Fig. 1D or Fig. 2D). So the tree can be virtually read from left to right or top to bottom in a consistent way. Several of these matrix-like elements can be combined without hampering analysis by visual overloading (Supplementary Fig. S5).

## 4 Discussion

To avoid some of the common problems associated scRNA-seq data visualization, we designed scBubbletree to convey key properties of cells in ways that are visually easily accessible and scalable. These properties include quantitative information about local and global similarity of transcription patterns and cell density. The properties can be flexibly linked to stacks of visually easily accessible additional information, e.g. on marker gene expression, cell type, or information from multiomics experiments such as immune cell receptors, surface-protein expression or methylation levels. The quantitative nature of its output lends scBubbletree to applications not only to research but also to quality control (Supplementary Section 2.4).

Yet, scBubbletree has clearly limitations. For instance, as a part of computing the dendrogram, the user has to choose a resolution. Although scBubbletree provides tools that support this choice, some familiarity with the underlying clustering concepts is required which precludes use of scBubbletree in a black-box manner. Furthermore, estimating the clustering resolution with *get_k* and *get_r* can be challenging in terms of computing times and memory for datasets of millions of cells, as these functions perform repetitive clustering simulations for a set of clustering resolutions. While the computational burden can be mitigated by parallelization, scBubbletree does not yet implement data representations that minimize memory requirements. In terms of its visualization capabilities, the current version of scBubbletree provides only the most essential functions for visualization of cell attributes. Future developments of the code will enrich its visualization repertoire in accordance with new developments in single-cell multiomics.

