## Supplementary material for "scBubbletree: Quantitative visualization of single cell RNA-seq data"

### scBubbletree: computational approach for visualization of single cell RNA-seq data — Supplementary material

This document includes:
- Supplementary Algorithm 1
- Supplementary Section 1-5
- Supplementary Figure 1-15

**Algorithm 1** Bubbletree generation.
Functions: min=minima of input values (function *min*, R-package base), sample=sampling with replacement (function *sample*, R-package base), HC=hierarchical clustering (function *hclust*, R-package stats), BBS=bootstrap branch support (function *prop.clades*, R-package ape)

**Input:**

$k$ ▷ number of clusters
$B$ ▷ number of bootstrap iterations
$C_1^{n_1 \times f}, C_2^{n_2 \times f}, \ldots, C_k^{n_k \times f}$ ▷ subsets of $A^{n \times f}$ for $k$ clusters
$n_1, n_2, \ldots, n_k$ ▷ number of cells in each cluster
$N_{eff}$ ▷ number of cells to draw from cluster

**Output:**

$\hat{H}$ ▷ consensus hierarchical dendrogram (bubbletree)
$H_b$ ▷ bootstrap hierarchical dendrograms

**for** $b = 1$ to $B$ **do**
  **for** $i = 1$ to $k - 1$ **do**
    $n_i' \leftarrow \min(n_i, N_{eff})$
    $X^{n_i' \times f} \leftarrow \text{sample}(n_i', C_i^{n_i \times f})$ ▷ draw $n_i'$ rows from $C_i^{n_i \times f}$
    **for** $j = i + 1$ to $k$ **do**
      $n_j' \leftarrow \min(n_j, N_{eff})$
      $Y^{n_j' \times f} \leftarrow \text{sample}(n_j', C_j^{n_j \times f})$ ▷ draw $n_j'$ rows from $C_j^{n_j \times f}$
      $D_b^{ij} \leftarrow \frac{1}{n_i' n_j'} \sum_{\substack{\forall \mathbf{x} \in X \\ \forall \mathbf{y} \in Y}} \sqrt{(\mathbf{x} - \mathbf{y})^2}$ ▷ mean inter-cluster Euclidean distance between cells (row vectors) of $X$ and $Y$
    **end for**
  **end for**
  $H_b \leftarrow \text{HC}(D_b^{ij})$ ▷ hierarchical clustering with average link
**end for**
$\hat{D}^{ij} \leftarrow \frac{1}{B} \sum_{b=1}^{B} D_b^{ij}$ ▷ inter-cluster distance for $i = 1, \ldots, k - 1$ and $j = i + 1, \ldots, k$
$\hat{H} \leftarrow \text{HC}(\hat{D}^{ij})$ ▷ consensus hierarchical clustering with average link
$\hat{H} \leftarrow \text{BBS}(\hat{H}, H_b)$ ▷ bootstrap branch support

### 1 Impact of clustering algorithm on biological cluster homogeneity

Quantitative comparison of the five clustering algorithms available as part of scBubbletree was performed with datasets A and B (Supplementary Section 2): k-means, Louvain (original, LMR, SMR), and, partially also Leiden.

k-means clustering was performed with $k = 5$ and $k = 24$ for dataset A and B, respectively. Clustering resolution parameter $r = 0.0025$ and $r = 0.79$ were used as input for the GCD methods for clustering of dataset A and B, respectively. Leiden clustering failed for dataset B due to the large size of the dataset.

The clustering results were compared with the adjusted RAND index (ARI) (function *adj.rand.index*, R-package fossil, version 0.4) [Hubert and Arabie, 1985]. ARI is a measure of similarity between two clustering results, yielding values between +1 (concordant results) and -1 (discordant results).

All five clustering methods produced similar results for dataset A (Supplementary Fig. S11A), and ARI was close to 1 between each pair of methods. The four GCD methods generated identical results (pairwise ARI $= 1$). For dataset B we observed considerable differences between the clustering results of the five methods (Supplementary Fig. S11C). The two Louvain variants, LMR and SMR, had similar results and ARI $= 0.93$. We also observed high degree of similarity (ARI $\approx$ 0.79) between the original Louvain method and LMR and SMR. ARIs were smaller (ARI $\in [0.66, 0.68]$) between the GCD methods and k-means.

The clustering results were also compared based on their weighted Gini impurities (WGIs). The WGIs were computed using the cell line labels of dataset A and the PBMC subtype labels from annotation set *l1* and *l2* of dataset B. For dataset A we saw WGIs close to 0 (low impurity) for all five clustering methods (Supplementary Fig. S11B). The GCD methods yielded similar WGIs, which were slightly lower than that of k-means. For dataset B, the GCD methods had systematically lower WGIs than k-means in terms of both annotation sets (Supplementary Fig. S11D). While the original Louvain algorithm marginally outperformed LMR and SMR with *l1*, application of SLM and LMR resulted in more homogeneous clusters than the original Louvain algorithm with *l2*.

In summary, we saw that the GCD methods tend to generate more homogenous clustering results than k-means, while the different Louvain variants and Leiden generated comparable results. Other clustering algorithms implemented in e.g. PhenoGraph [Levine et al., 2015] or TooManyCells [Schwartz et al., 2020], were not included in the benchmarking analysis as they rely on GCD method for clustering and are likely produce comparable results as the GCD methods implemented in Seurat.

### 2 Datasets and data processing

#### 2.1 Dataset A

scRNA-seq dataset A contains a mixture of 3,918 cells from five human lung adenocarcinoma cell lines (HCC827, H1975, A549, H838 and H2228). This dataset has been generated for the purposes of benchmarking of pipelines and methods for scRNA-seq analysis [Tian et al., 2019]. Sample collection and data processing details are provided in the respective publication [Tian et al., 2019]. Key data processing steps are summarized in the following. The single cell library were prepared using 10x Chromium platform and sequenced with Illumina NextSeq 500. Raw data was processed with Cellranger and analyzed by scPipe. demuxlet [Kang et al., 2018] was used to predict the most probable identity of each cell line based on the known genetic differences between the different cell lines. We obtained the dataset (with code *sc_10x_5cl*) and the cell type predictions from https://github.com/LuyiTian/sc_mixology.

#### 2.2 Dataset B

scRNA-seq dataset B contains transcriptome data of 161,764 peripheral blood mononuclear cells (PBMCs) from eight healthy volunteers enrolled in a HIV vaccine trial [Hao et al., 2021]. Sample collection and data processing details are provided in the publication [Hao et al., 2021] and are summarized in the following. Samples were collected at three time points: immediately before (day 0), 3 days, and 7 days following administration of a HIV vaccine. All samples have been profiled using 10x Chromium protocol, followed by Illumina NovaSeq 6000 sequencing. Alongside the single cell transcriptomes, CITE-seq technology was used with up to 228 antibodies to profile the expression of cell-surface proteins. This multimodal data was processed with a weighted nearest neighbor (WNN) procedure to identify different cellular states in PBMCs. Cell types were predicted by this approach for each cell at two levels of resolution: *l1* with 8 cell types and *l2* with 31 cell types. The predictions are available as part of the meta data associated with the raw scRNA-seq data.

#### 2.3 Data processing

Data processing of dataset A and B was performed with R-package Seurat (version 4.1.0). Normalization of gene expressions was done with the function *SCTransform* [Hafemeister and Satija, 2019] using default parameters, and principal component analysis (PCA) was performed with function *RunPCA* based on the 5,000 most variable genes in the dataset identified with the function *FindVariable-*

*Features*. In both datasets we saw that the first 15 principal components capture most of the variance in the data, and the proportion of variance explained by each subsequent principal component was negligible (Supplementary Fig. S1A-B). Thus, we used the single cell projections (embeddings) in 15-dimensional feature space, $A^{3,918\times15}$ and $A^{161,764\times15}$, as input of scBubbletree. Dimensionality reduction of $A^{3,918\times15}$ and $A^{161,764\times15}$ to 2-dimensions (2D) by UMAP and t-SNE was performed with Seurat's functions *RunUMAP* and *RunTSNE*, respectively, using default parameters.

#### 2.4 Quality control with scBubbletree

The quantitative nature of its output lends scBubbletree to applications not only to research but also to quality control. In the following we demonstrate this by quality control checks of datasets A and B.

##### 2.4.1 Dataset A

The quality control of dataset A is summarized in the respective study [Tian et al., 2019]. We used scBubbletree to evaluate the relative frequency of mitochondrial RNA in the different bubbles (Supplementary Fig. S12A), which is an indicator of poor sample quality associated with high fraction of apoptotic or lysing cells. On average, about 19% percent of the RNA molecules originated from mitochondrial genes. We also observed bimodal distributions of the number of detected genes and RNA molecules in bubbles 0, 1 and 2 (Supplementary Fig. S12B-C). Nevertheless, the numbers of detected genes and RNA molecules were sufficiently high for downstream analysis.

##### 2.4.2 Dataset B

Dataset B contains PBMC samples from 8 donors at 3 time points. Using donor and time used as categorical cell features, we visualized the relative frequency of cells from a specific donor and time across the different bubbles (Supplementary Fig. S13). This allowed us to check for compositional biases, which is essential to e.g. identify novel cell types (e.g. tumor cells), or to detect systematic biases (e.g. batch effects) in a subset of the samples. This analysis revealed compositional biases in the samples of specific donors, for instance, all three samples of donors P2 and P3 had relatively higher frequency of bubble 0 and 2 cells, respectively. To conduct a quantitative differential abundance analysis of cell type compositions [Buettner et al., 2021], scBubbletree provides tables with cell counts for each pair of sample and bubble.

As part of the quality checks we also looked for bubbles of cells with unusually high relative frequency of mitochondrial RNA (Supplementary Fig. S14A). We saw similar distributions of mitochondrial gene expression across the different bubbles, with about 95% of the cells having mitochondrial gene expression lower than 10%. In contrast to this, we saw differences in number of detected genes in the cells of the different bubbles. For instance, the cells in bubble 18 had lower numbers of detected genes (median number of genes $< 1{,}000$) and RNA molecules (median number of RNA molecules $< 2{,}500$) (Supplementary Fig. S14B-C). This may be explained by the fact that most cells in bubble 18 are classified as platelets. This cell type is devoid of nucleus and genomic DNA, and hence is incapable of de novo transcription [Melchinger et al., 2019].

### 3 Gene expression of human cancer cell lines

We obtained gene expression data for 1,019 human cancer cell lines from the Cancer Cell Line Encyclopedia [Barretina et al., 2012] (https://www.ebi.ac.uk/gxa/-experiments/E-MTAB-2770/Results, obtained on 20.04.2022), and selected among them 69 cell lines that contained the keyword “lung adenocarcinoma” in their names. Expressions (transcript per million; TPM) were available for 56,443 genes. To quantify the distance in gene expression between a pair of cell lines, we computed the Euclidean distance between the corresponding gene expression vectors (function *dist*, R-package stats, version 4.2). This procedure was repeated for 2,346 pairs of cell lines. In the distribution of Euclidean distances between the cell lines we saw a conspicuously small distance between cell lines H1975 and HCC827 (Supplementary Fig. S3). The distances between the remaining cell lines were closer to the mean of the distribution.

### 4 Evaluation of the bubbletree of dataset B based on annotation set *l2*

The composition of the bubbles was also investigated based on the more detailed annotation set *l2*. Using the function *get_cat_tiles* we visualized the within-bubble relative frequencies of 31 PBMC subtypes from *l2* (heatmap in Supplementary Fig. S5A) and made the following observations.

First, the bubbles of clade ‘b’ were enriched with cells from distinct populations of monocytes and DCs (Supplementary Fig. S5A). Bubbles 1, 3 and 7, which mostly contained $CD14^{+}$ classical monocytes, were characterized by high CD14 and low CD16 protein and RNA expression (RNA heatmap in Supplementary Fig. S5B). About 77% of all monocytes were found in these bubbles (Supplementary Fig. S15), and this was comparable to the expected relative frequency of classical monocytes in humans [Patel et al., 2017]. Bubbles 13 and 19 contained a mixture of $CD14^+$ classical monocytes and $CD16^+$ non-classical monocytes. As CD14 and CD16 were both highly expressed in bubble 13 and bubble 19 cells, it is likely that these bubble are enriched with intermediate monocytes. Interestingly, bubble 19 was also enriched with cells characterized as doublets. Finally, bubble 12 was enriched with $CD16^+$ non-classical monocytes, whereas bubble 15 contained a mixture of $CD14^+$ classical monocytes and three populations of DCs: conventional DC1 and DC2 (cDC1 and cDC2) and $Axl^+$ $Siglec6^+$ DC (ASDC).

Second, the bubbles of the B cell subclade were enriched with functionally distinct subtypes of B cell, i.e. bubble 10 contained mainly naive B cells (98% of the bubble cells), bubble 11 contained a mixture of intermediate (39% of the bubble cells), memory (55% of the bubble cells) and a small fraction of naive B cells (5% of the bubble cells), while bubble 22 was enriched with plasmablasts (98% of the bubble cells). Bubbles 2 and 16 made up the NK subclade and were enriched with cells from two functionally different NK subtypes that are characterized by distinct expression of the surface marker CD56 (ADT heatmap in Supplementary Fig. S5B): $CD56^{bright}$ and $CD56^{dim}$. In bubble 2 we saw an enrichment of $CD56^{dim}$ NK cells (99% of the bubble cells), and bubble 16 contained a mixture of $CD56^{dim}$ and $CD56^{bright}$ NK cells (67% and 29% of the bubble cells, respectively).

Third, the remaining bubbles of clade 'a' were enriched with cells from different T cell subtypes. Functionally and/or developmentally related subtypes of T cells were found in common subclades, for instance, bubble 5 was enriched with naive $CD8^+$ T cells (98% of the bubble cells), while bubble 0 was enriched with naive $CD4^+$ T cells (87% of the bubble cells). These two bubbles accounted for about 18% of all cells in the sample and about 50% of all T cells. Similarly, bubble 4 and 6 were primarily enriched with different memory subtypes of $CD4^+$ and $CD8^+$ T cells. Bubbles 9 and 14 were enriched with $CD8^+$ memory T cells, mucosal associated invariant (MAIT) T cells and $\gamma\delta$ T cells, whereas bubbles 8 and 17 were enriched with T cells which share phenotype with NK cells, including $\gamma\delta$ T cells, $CD4^+$ T cells with cytotoxic activity (CD4 CTL) and $CD8^+$ effector memory T cells (CD8 TEM). The small bubbles 21 and 23 contained proliferating lymphocytes and hematopoietic stem and progenitor cell (HSPC), respectively.

Fourth, the outgroup bubble 18 was enriched with platelets (99% of cells in that bubble), whereas bubble 20 contained plasmacytoid dendritic cells (pDCs, 96% of cells in bubble) and a small fraction of ASDCs (4% of the cells in bubble).

### 5 Quantifying uncertainty with Highest Density Intervals (HDIs)

The HDI was used to summarize the most credible points of a distribution (function *hdi*, R-package bayestestR, version 0.13). All points within the HDI have a higher probability density than points outside the interval, and the total mass of points inside the 95% HDI is 95% of the distribution.

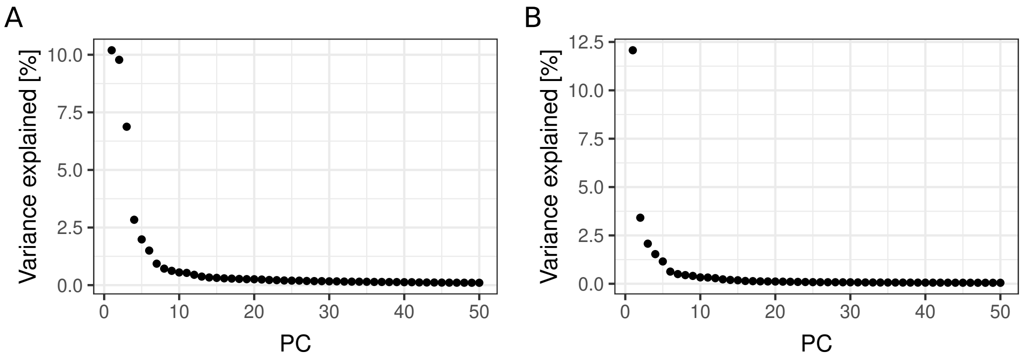


Supplementary Figure S1. Percent of variance explained by principal components 1 to 50 in data set A (panel A) and B (panel B).

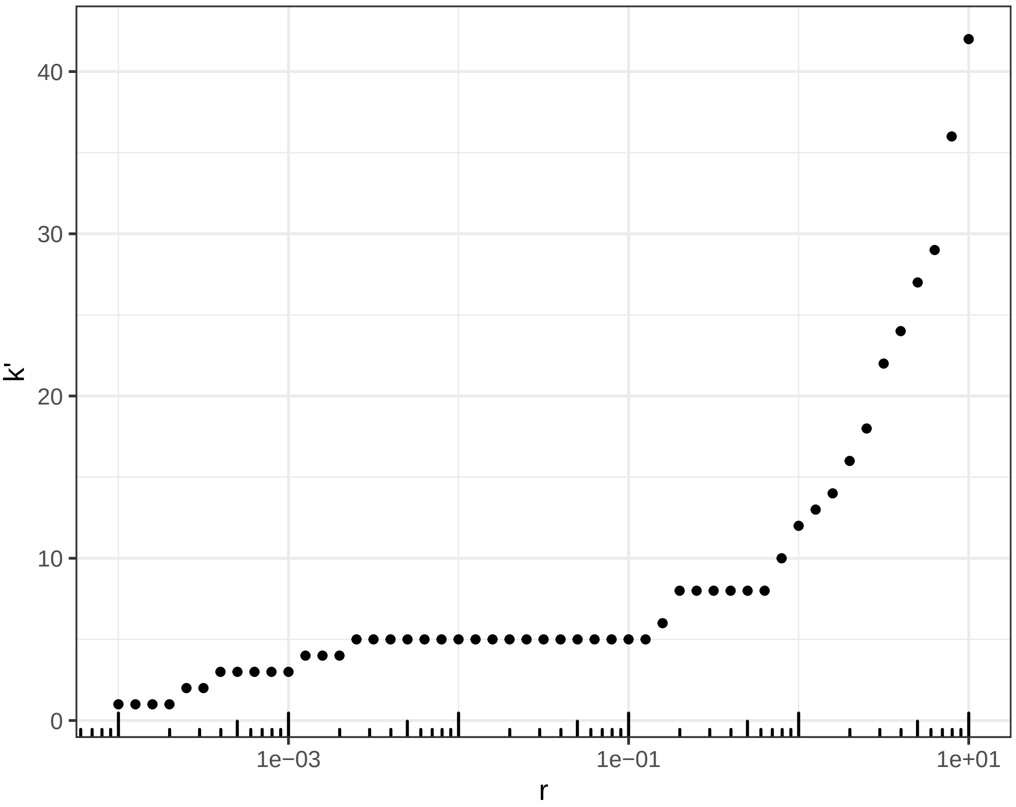


Supplementary Figure S2. Relationship between resolution ($r$) and the number of identified communities ($k'$) by Louvain custering of dataset A.

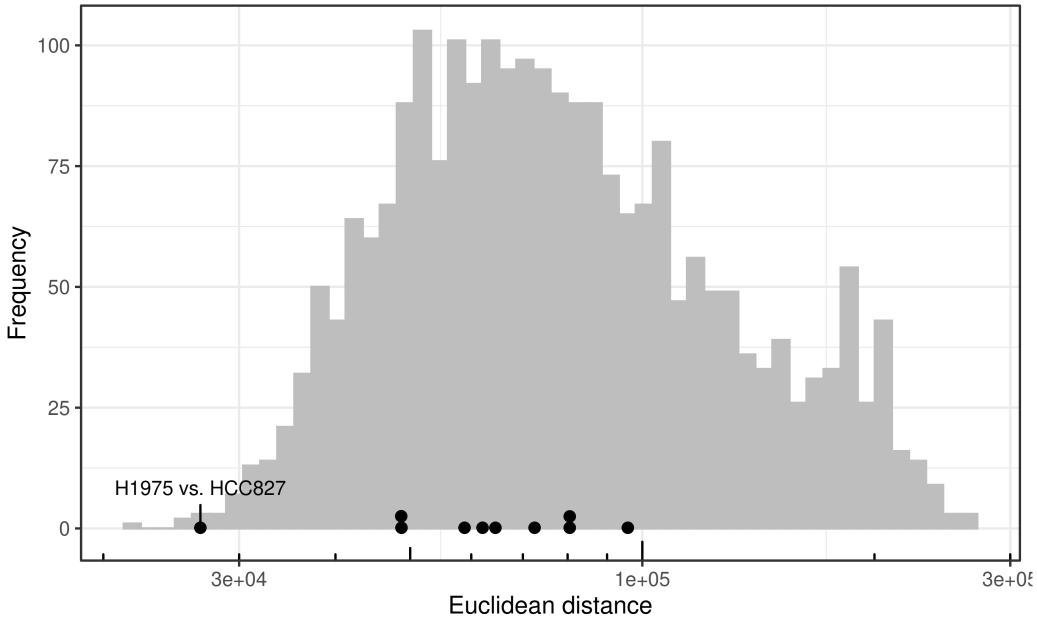


Supplementary Figure S3. Histogram of distances in gene expression between 2,346 pairs of 69 human lung adenocarcinoma cell lines. Distances between cell lines H838, H2228, A549, H1975 and HCC827 are shown with black filled points. The distance between cell lines H1975 and HCC827 is annotated with a black label. Vertical jitter was added to avoid overplotting of black points.

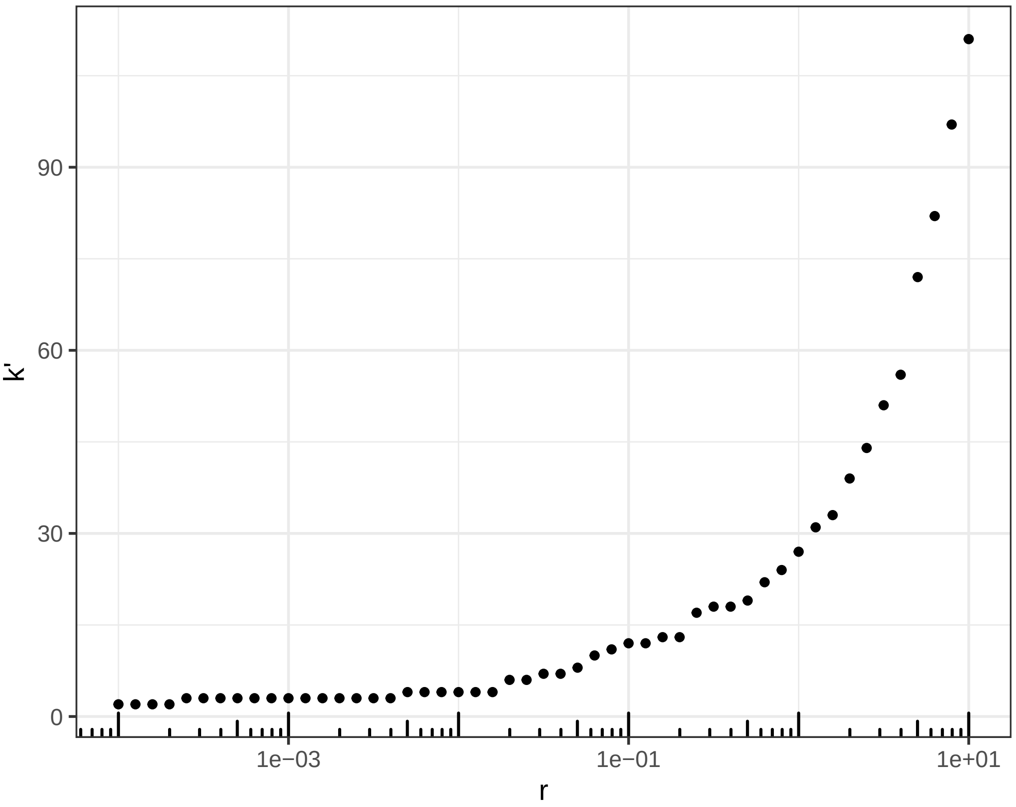


Supplementary Figure S4. Relationship between resolution ($r$) and the number of identified communities ($k'$) by Louvain custering of dataset B.

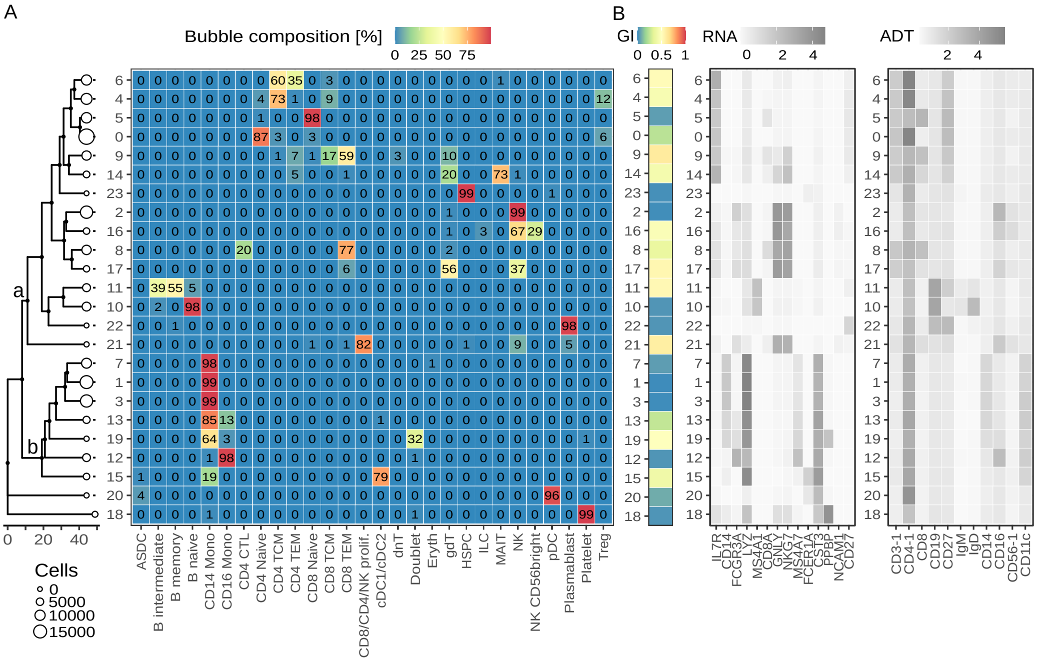


Supplementary Figure S5. Cell type and bubble composition of dataset B and marker gene/protein expression. (A) Relative frequencies (%) of cell types (*l2* annotation set; x-axis) across bubbles (y-axis) rounded to the nearest integer. Rows integrate to 100% (up to rounding error). A simplified version of the bubbletree from Fig. 2C is appended to panel A. For improved readability proliferating CD8 T, CD4 T and NK cells are jointly shown under the label CD8/CD4/NK prolif. cDC1 and cDC2 cells are jointly shown under the label cDC1/cDC2. (B) Three heatmaps with: bubble-specific Gini impurity (GI) computed based on *l2* labels (left panel of B); mean normalized expression (RNA) of 14 genes (x-axis) in each bubble (y-axis) (middle panel of B); and mean normalized expression (ADT) of 11 surface-proteins (x-axis) in each bubble (y-axis) (right panel of B).

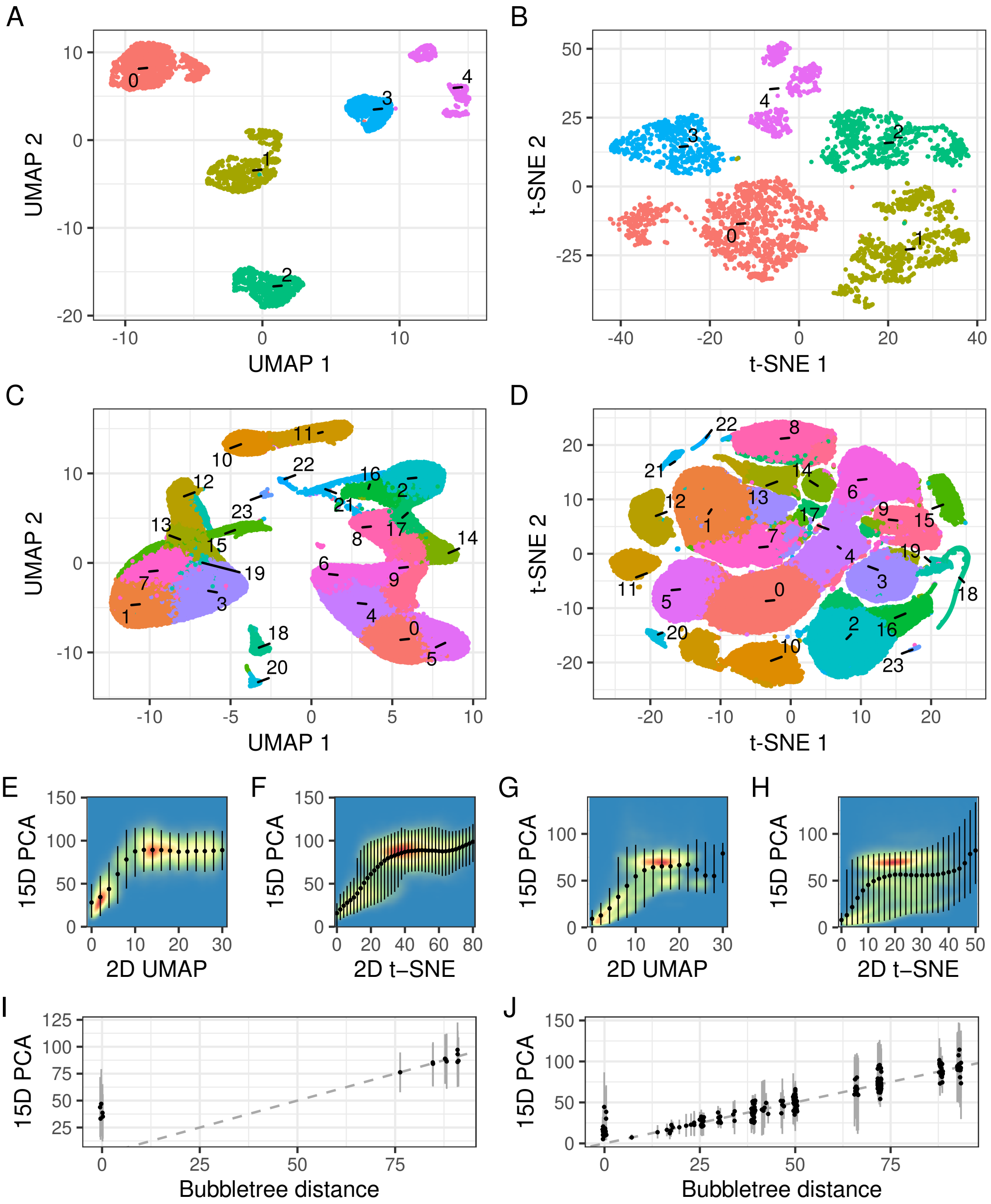


Supplementary Figure S6. 2D UMAP and t-SNE visualization and distance structure of dataset A and B. (A, B) 2D UMAP and t-SNE of dataset A. (C, D) 2D UMAP and t-SNE of dataset B. Filled points represent cells, color coded according to their bubble IDs. (E, F, G, H) 2D density of distances in 2D UMAP and t-SNE vs. 15D PCA space between $10^6$ cell pairs, randomly selected with replacement from data sets A (panels E, F) and B (panels G, H). Blue, yellow and red density gradient color codes for low, medium and high density, respectively. Points are means and vertical error bars are 95% highest density intervals (HDIs, Supplementary Section 5) of the cell-cell distances in 15D PCA space, computed over the data of overlapping x-axis bins obtained by sliding window approach (sliding step = 2, window length = 4). (I, J) Distances between bubbles in the bubbletree (x-axis, with random jitter of points and error bars along the horizontal axis to reduce overplotting) and average Euclidean distances between cells in 15D PCA space (y-axis) for dataset A (panel I) and B (panel J). Error bars are 95% HDIs of the distances in 15D PCA space between $10^4$ pairs of cells randomly selected with replacement from each pair of bubbles. Dashed gray lines are diagonals.

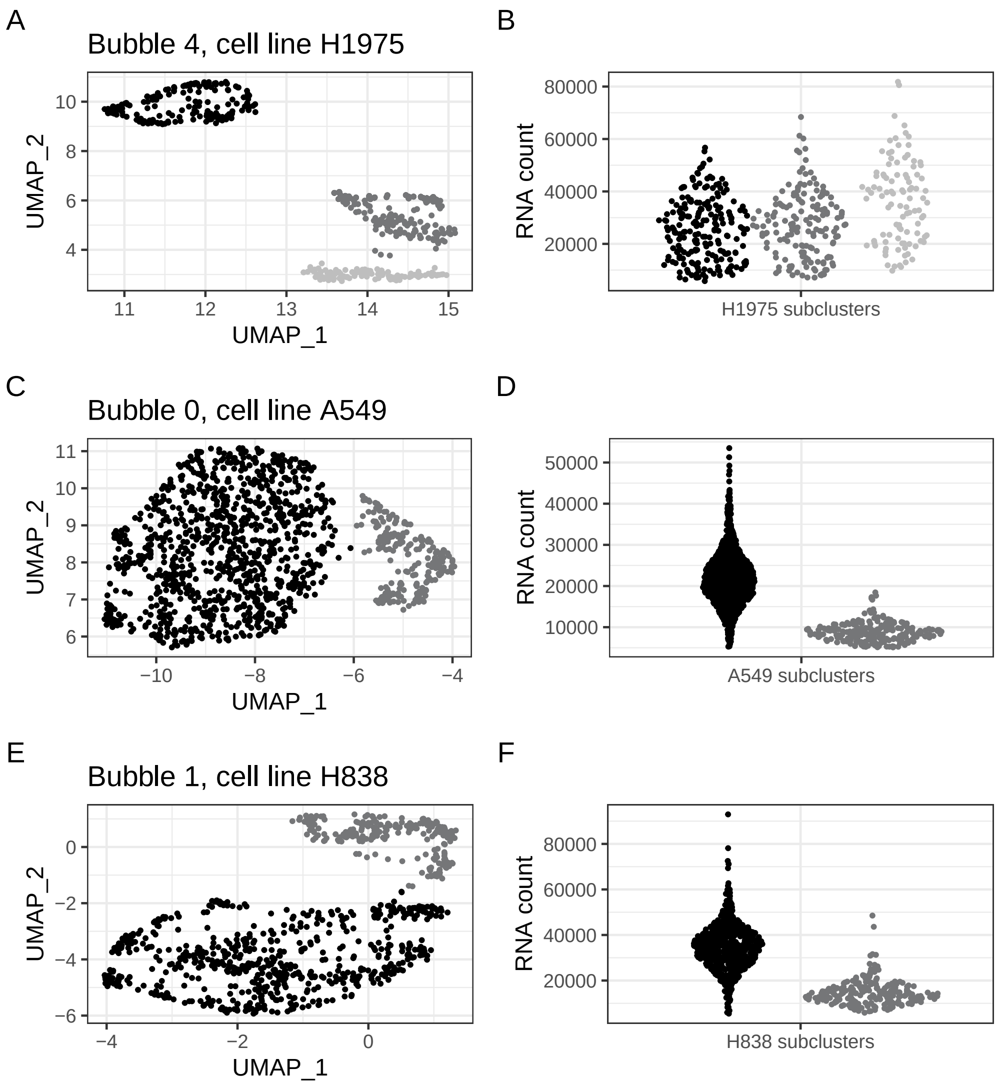


Supplementary Figure S7. Cell line subclusters in 2D UMAP space of dataset A. (A, C, E) Cells in different subclusters (indicated by different shades of gray) in bubble 4 (panel A), bubble 0 (panel C) and bubble 1 (panel E). Cells from bubble 4, 0 and 1 are enriched with cells from cell line H1975, A549 and H838, respectively. (B, D, F) Distribution of the number of RNA molecules detected in cells from each subcluster of bubble 4 (panel B), 0 (panel D) and 1 (panel F).

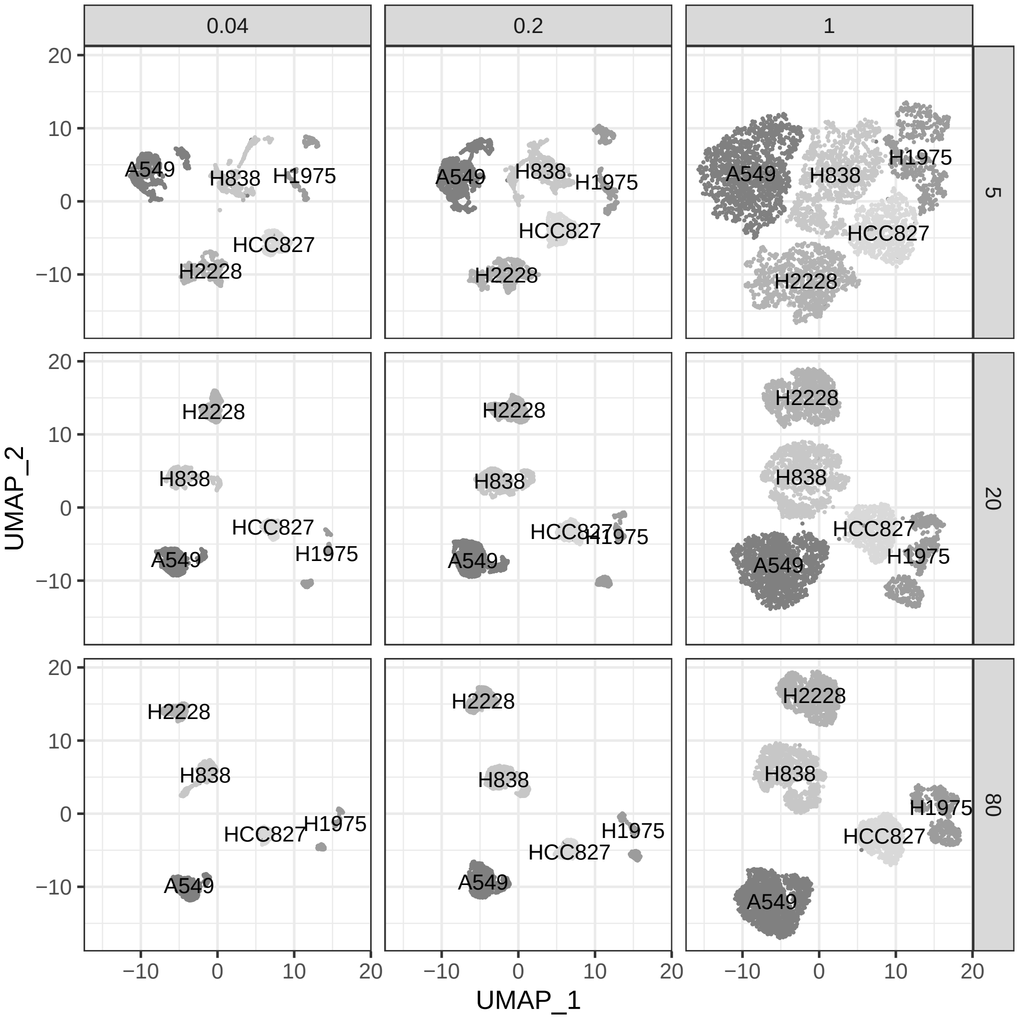


Supplementary Figure S8. Effects of UMAP hyperparameters *min_dist* and *n_neighbors* on the 2D UMAP embeddings of dataset A. Filled points are cells, gray-level coded according to their cell lines. Column panels correspond to *min_dist*=0.04, 0.02 and 1; row panels correspond to *n_neighbors*=5, 20 and 80. Default values were used for the remaining two UMAP hyperparameters, *d*=2 and *n_epoch*=300. UMAP dimensionality reduction was performed with the function *RunUMAP* from the R-package Seurat. Visualization was performed with the R-package ggplot2. Parameter legend: *min_dist*: the neighborhood size to use for local metric approximation; *n_neighbors*: parameter for controlling the layout; *n_epoch*: the number of training epochs to be used in optimizing the low dimensional embedding; *d*: the number of dimensions.

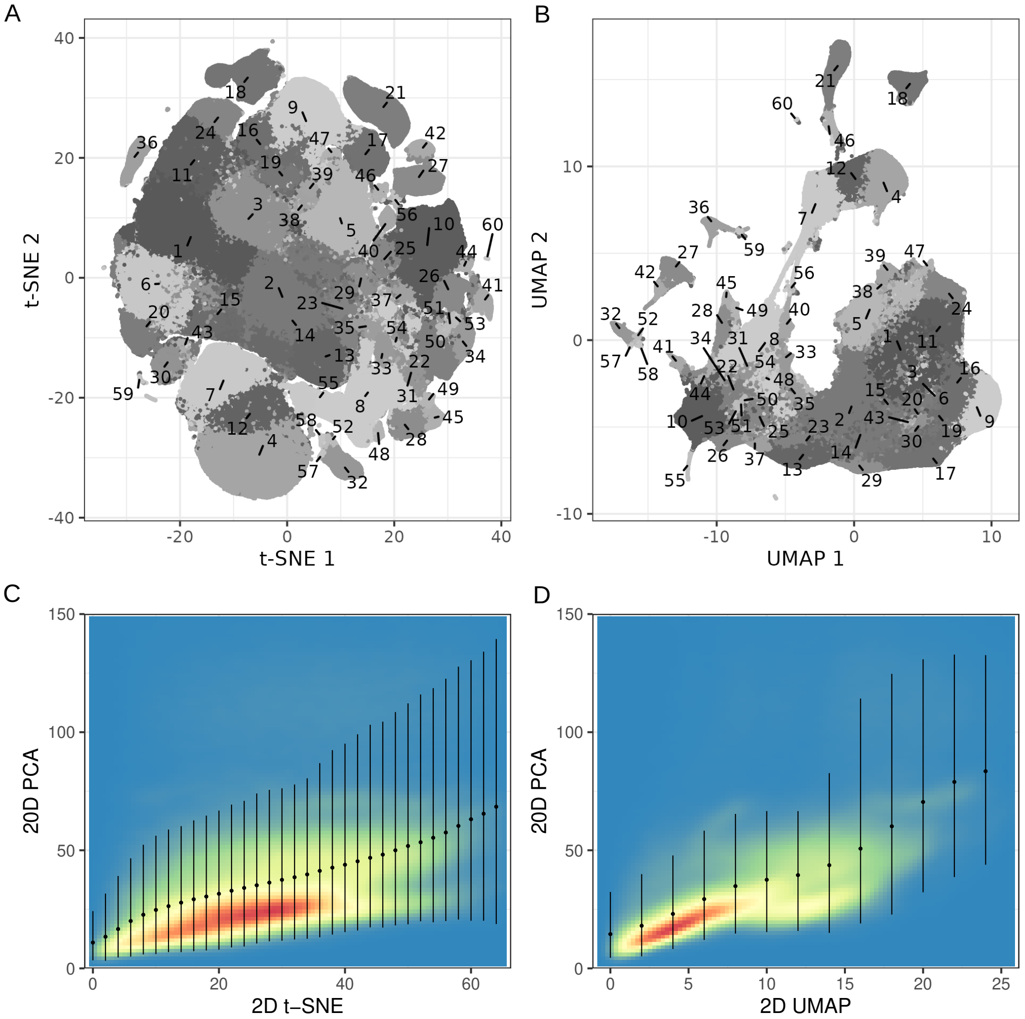


Supplementary Figure S9. 2D t-SNE and UMAP and distance structure of scRNA-seq data of 1.3 million mouse brain cells. (A) 2D t-SNE plot. Filled dots are cells, grey-level coded according to their cluster IDs. Labels show cluster IDs. PCA, t-SNE and graph-based clustering data were downloaded from https://support.10xgenomics.com/single-cell-gene-expression/datasets/1.3.0/1M_neurons? on 26.10.2022. (B) UMAP plot generated with function *umap* (R-package umap, version 0.2.10). Filled dots are cells color coded according to their cluster IDs. Labels show cluster IDs. (C, D) 2D density of distances between cells in 2D t-SNE (panel C) and 2D UMAP (panel D) vs. 20D PCA space between $10^6$ cell pairs randomly selected with replacement. Blue to red gradient colors reflect low to high density. Points are means and vertical error bars are 95% highest density intervals (HDIs, Supplementary Section 5) of the cell-cell distances in 20D PCA space, computed over the data of overlapping x-axis bins obtained by sliding window approach (sliding step=2, window length=4).

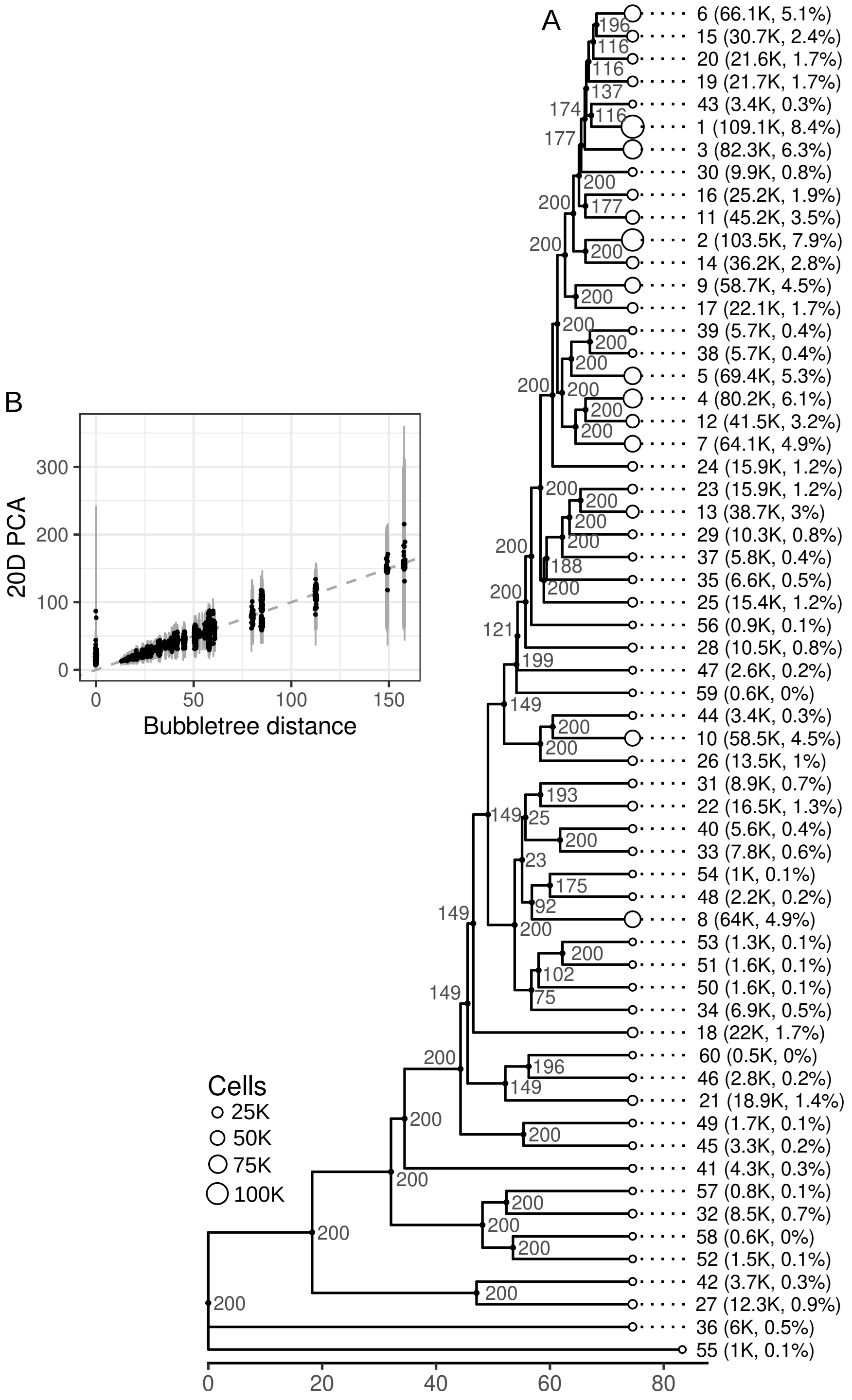


Supplementary Figure S10. bubbletree of scRNA-seq data of 1.3 million mouse brain cells. (A) Bubbletree generated with function *get_bubbletree_dummy* based on 20D PCA data and cluster assignments. We performed $B = 200$ bootstrap iterations and drew samples with up to $N_{eff} = 400$ cells from each cluster to estimate inter-cluster distances. PCA and graph-based clustering data were downloaded from https://support.10xgenomics.com/single-cell-gene-expression/datasets/1.3.0/1M_neurons? on 26.10.2022. (B) Bubbletree distances (x-axis, with random jitter along the horizontal axis to reduce overplotting) and mean Euclidean distances between cells in 20D PCA space (y-axis). Error bars are 95% highest density intervals (HDIs, Supplementary Section 5) of the distances in 20D PCA space between $10^4$ pairs of cells randomly selected with replacement from each pair of bubbles. Diagonal is shown as dashed gray line.

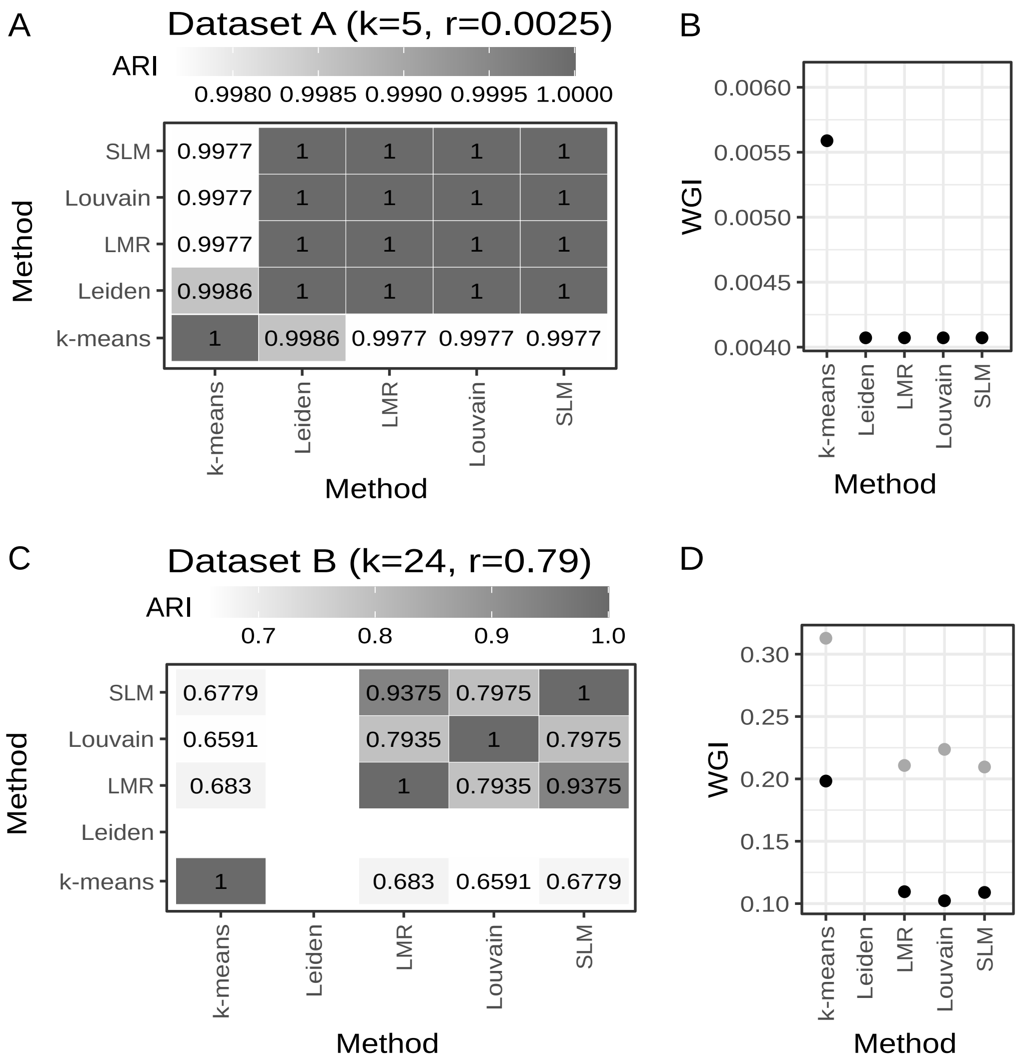


Supplementary Figure S11. Comparison of clustering algorithms. (A, C) Tiles are gray-level coded and labeled according to the adjusted RAND index (ARI) between pairs of clustering solutions from five clustering algorithms implemented in scBubbletree. ARIs are shown for dataset A (panel A) and dataset B (panel C). Clustering resolution parameter $k = 5$ and $k = 24$ were used as input for k-means for clustering of dataset A and B, respectively. Clustering resolution parameter $r = 0.0025$ and $r = 0.79$ were used as input for the graph-based community detection methods for clustering of dataset A and B, respectively. Leiden clustering failed for dataset B due to the large size of the dataset. (B) Points are weighted Gini impurity (WGI) indices for the five clustering algorithms and the cell line labels of dataset A. (D) Points are WGIs for the five clustering algorithms and the PBMC subtype labels from annotation set *l1* and *l2* of dataset B. Louvain: original Louvain algorithm, LMR: Louvain algorithm with multilevel refinement, SLM: smart local moving, Leiden: Leiden algorithm.

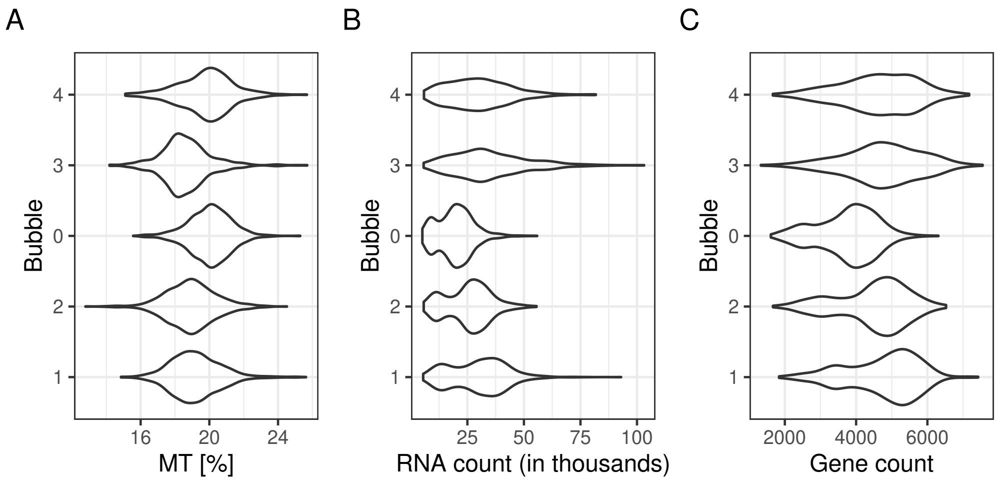


Supplementary Figure S12. Quality control of dataset A. Violins show in different bubbles (y-axis) the distribution of (A) the relative contribution to the total gene expression in a cell coming from mitochondrial genes, (B) the number of detected RNA molecules and (C) the number of detected genes.

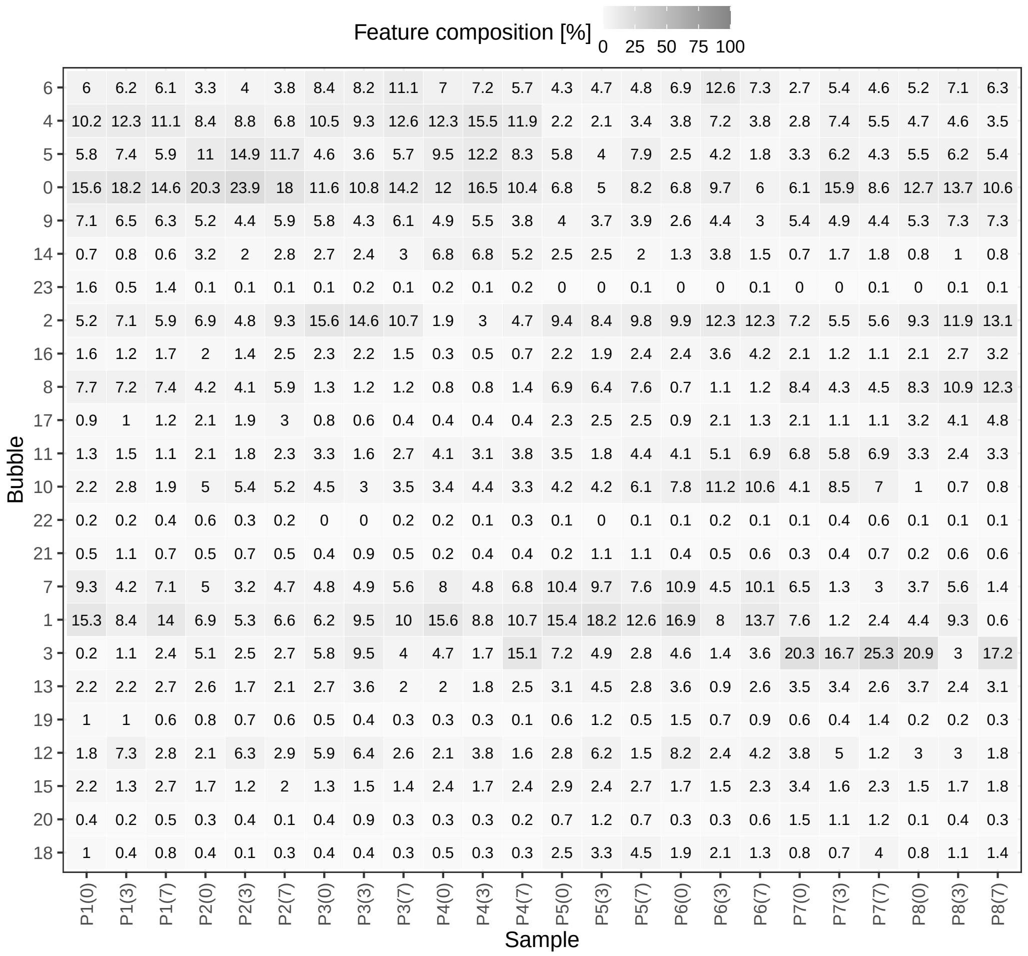


Supplementary Figure S13. Compositional analysis of dataset B. Tile labels in the heatmap are relative frequencies of cells coming from a specific sample (x-axis) across the different bubbles (y-axis). Columns integrate to 100%. Samples come from a combination of donor (P1-P8) and time points (0, 3 and 7). Relative frequencies are rounded to the nearest tenth.

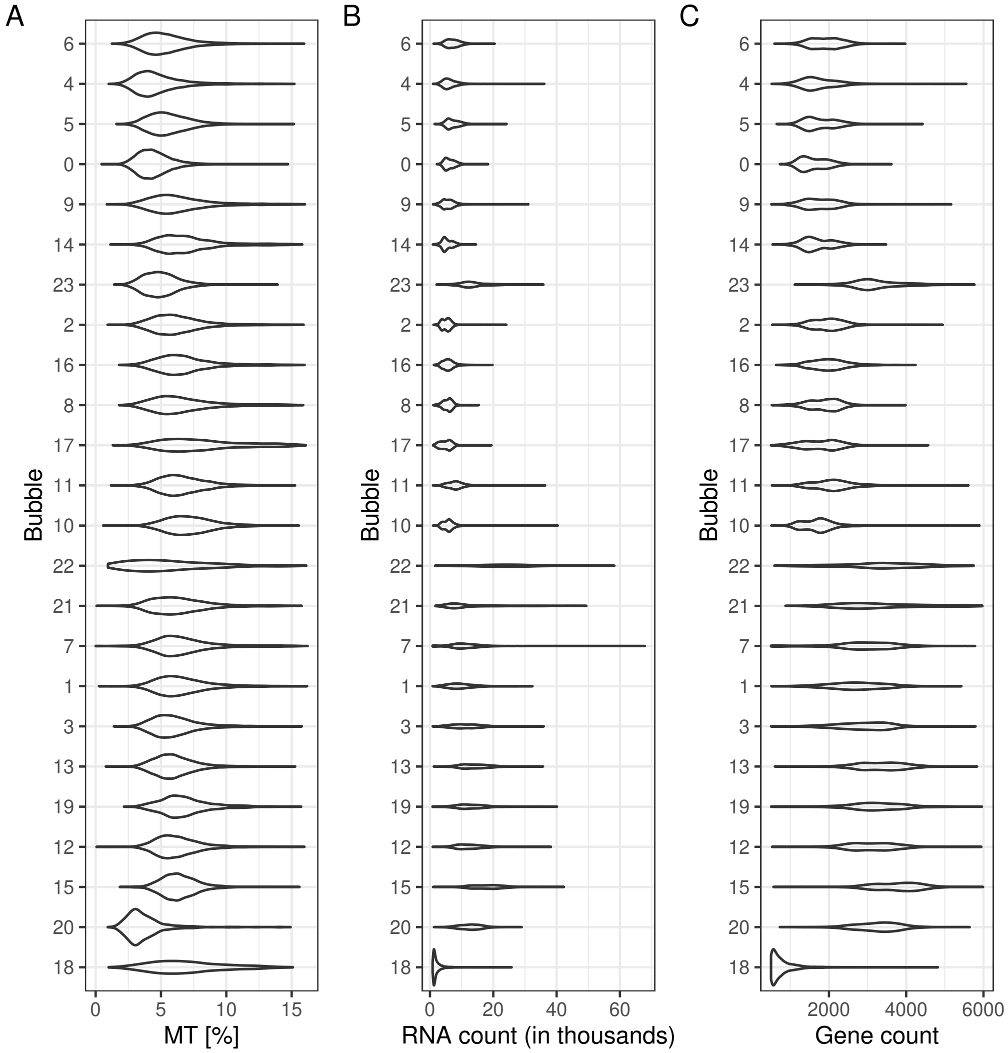


Supplementary Figure S14. Quality control of dataset B. Violins show in different bubbles (y-axis) the distribution of (A) the relative contribution to the total gene expression in a cell coming from mitochondrial genes, (B) the number of detected RNA molecules, and (C) the number of detected genes.

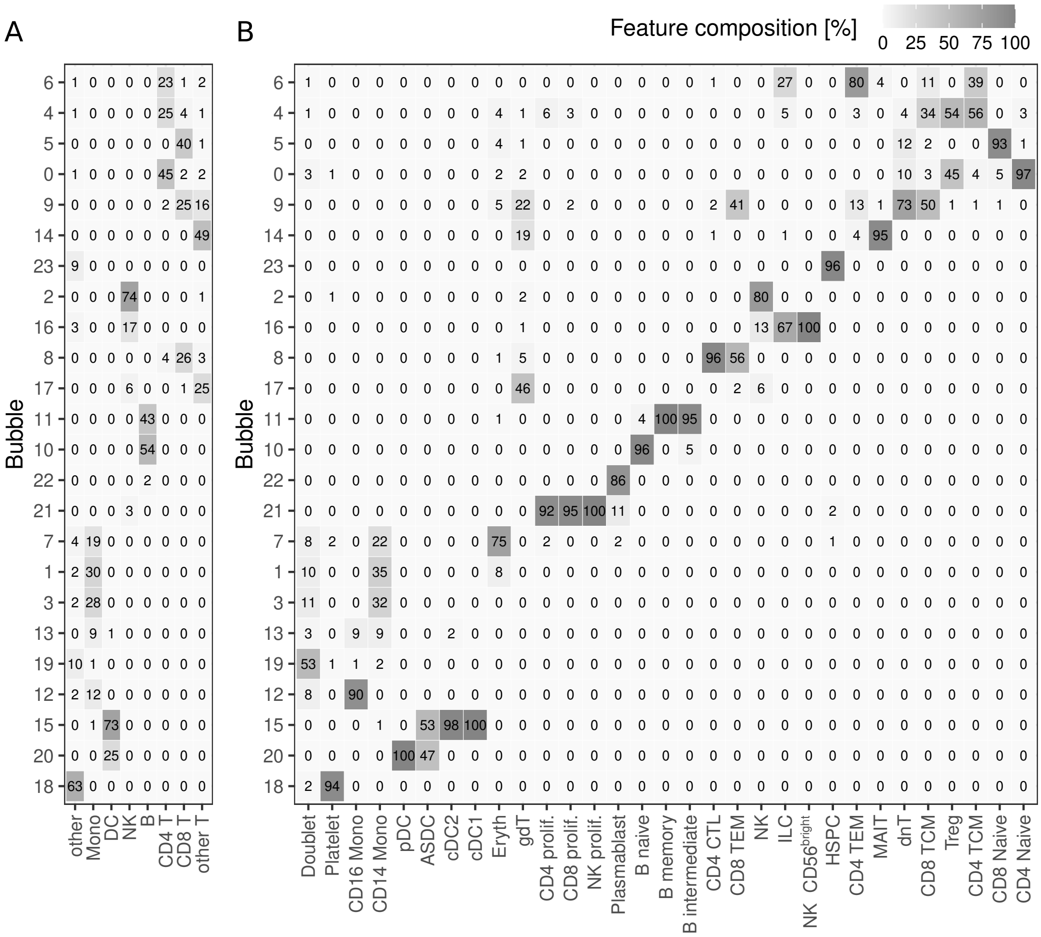

Supplementary Figure S15. Cell type composition across bubbles. Relative frequencies (%) of cell type labels (x-axis) from annotation set *l1* (panel A) and *l2* (panel B) across bubbles (y-axis). Relative frequencies are rounded to the nearest integer. Columns integrate to 100% (up to rounding error).
